# Altering the substitution and crosslinking of glucuronoarabinoxylans affects cell wall porosity and assembly in *Brachypodium distachyon*

**DOI:** 10.1101/2023.08.17.553603

**Authors:** Theodora Tryfona, Yanina Pankratova, Deborah Petrik, Diego Rebaque Moran, Raymond Wightman, Alberto Echevarria Poza, Xiaolan Yu, Parveen Kumar Deralia, Francisco Vilaplana, Charles T. Anderson, Mei Hong, Paul Dupree

## Abstract

- The Poaceae family of plants provides cereal crops that are critical for human and animal nutrition and also they are an important source of biomass. Interacting plant cell wall components give rise to recalcitrance to digestion, thus understanding the wall molecular architecture is important to improve biomass properties. Xylan is the main hemicellulose in grass cell walls. Recently, we reported structural variation in grass xylans, suggesting functional specialisation and distinct interactions with cellulose and lignin. Here, we investigated the functions of these xylans by perturbing the biosynthesis of specific xylan types.
- We generated CRISPR/Cas9 knockout mutants in *Brachypodium distachyon XAX1* and *GUX2* genes involved in xylan biosynthesis. Using carbohydrate gel electrophoresis we identified biochemical changes in different xylan types. Saccharification, cryo-SEM, subcritical water extraction and ssNMR were used to study wall architecture.
- *Bd*XAX1A and *Bd*GUX2 enzymes modify different types of grass xylan. *Brachypodium* mutant walls are more porous, suggesting the xylan substitutions directed by both *Bd*XAX1A andGUX2 enzymes influence xylan-xylan and/or xylan-lignin interactions.
- Since xylan substitutions influence wall architecture and digestibility, our findings open new avenues to improve cereals for food and to use grass biomass for feed and the production of bioenergy and biomaterials.

## Introduction

Most of the energy and renewable materials in plant lignocellulosic biomass is locked within secondary cell walls in cellulose and xylan in a dense matrix with lignin (McCann & Carpita, 2008). Grasses are considered an abundant renewable source of biomass for bioenergy and biomaterials production due to their year-round production and ability to grow on less fertile land and in semi-arid environments. Bamboo is a grass widely used as a particularly strong material with properties in some ways superior to wood, reflecting the remarkable grass cell wall strength (Shah *et al*., 2017). However, grass lignocellulosic biomass is intrinsically recalcitrant to enzymatic degradation, which hampers its use for biochemical production of biofuel.

The degree, type and distribution of substitutions on xylan are major factors affecting cell wall recalcitrance by forming cross-links with other xylan molecules and lignin (Grabber *et al*., 2000; Nishimura *et al*., 2018; Feijao *et al*., 2022; Nishimura *et al*., 2022), by influencing interactions with cellulose (Grantham *et al*., 2017) and by impairing the action of xylan hydrolytic enzymes (Biely *et al*., 2016). Xylan comprises a linear backbone of *β*-1,4-xylose (Xyl) residues commonly substituted with *α*-1,2-linked (4-*O*-methyl)-glucuronic acid (GlcA), *α*-1,3-linked arabinofuranose (Ara*f*) and acetylation at the *O*-2 and/or *O*-3 positions (Ebringerová & Heinze, 2000). In eudicots, GlcA and acetyl substitutions are positioned mostly on alternate Xyl residues of the backbone (Bromley *et al*., 2013; Busse-Wicher *et al*., 2014). It is this even pattern of decorations that allows xylan to adopt a twofold helical screw conformation, with the substitutions being oriented on one side of the xylan chain, putatively permitting xylan to form hydrogen bonds and stacking interactions with hydrophilic surfaces of cellulose. Randomly decorated xylan, on the other hand, adopts a flexible threefold conformation and therefore does not interact with cellulose hydrophilic surfaces in the same way (Grantham *et al*., 2017). Even spacing of xylan decorations is a highly conserved feature not only in Eudicots and Gymnosperms (Busse-Wicher *et al*., 2016; Martínez-Abad *et al*., 2017; Pereira *et al*., 2017) but also in grasses (Tryfona *et al*., 2023). Recently we proposed that grass cell wall xylan is structurally diverse, consisting of at least three different xylan domains namely <u>A</u>rabino<u>x</u>ylan with <u>e</u>ven distribution of Ara*f* substitutions (AXe); <u>G</u>lucurono<u>a</u>rabino<u>x</u>ylan with <u>c</u>lustered GlcA substitutions (GAXc) and <u>h</u>ighly <u>s</u>ubstituted <u>g</u>lucurono<u>a</u>rabino<u>x</u>ylan (hsGAX) (Tryfona *et al*., 2023). We hypothesised that these distinct xylan types might adopt different conformations and have different interactions with other wall components (Tryfona *et al*., 2023). Indeed, our recent ssNMR studies of never-dried *Brachypodium distachyon* tissues confirms that some xylan in grasses binds to cellulose in the two-fold conformation (Duan *et al*., 2021). Lignin-polysaccharide interactions in secondary cell walls of maize revealed by ssNMR were also suggested to be seen preferentially on xylan with three-fold conformation (Kang *et al*., 2019). Nevertheless, it is still unknown how the different xylan types are important in grass cell wall assembly.

Grass xylans have unique structural features. Although the substitutions consist mainly of Ara*f* and also GlcA substitutions similar to other plants (Tryfona *et al*., 2019), there are additionally, hydroxycinnamic acid groups (for example, feruloyl (Fer)- or coumaryl (Co) group) esterified to the *O*-5 position of Ara*f* residues (Mueller-Harvey *et al*., 1986; Hatfield *et al*., 2017; Feijao *et al*., 2022), and also Xyl*p*-Ara*f* substitutions (Wende & Fry, 1997; Tryfona *et al*., 2019; Zhong *et al*., 2022). Ferulation of grass xylan facilitates its dimerization or crosslinking to lignin via diferulate bridge formation with coniferyl or sinapyl lignin units (Hatfield *et al*., 1999; Hatfield *et al*., 2017), possibly increasing the strength and recalcitrance of the wall (Buanafina *et al*., 2010; Ralph, 2010; de Oliveira *et al*., 2015; de Souza *et al*., 2019). A covalent linkage between GlcA of glucuronoxylan and lignin has been reported in eudicots (Imamura *et al*., 1994; Nishimura *et al*., 2022). Covalent links between the GlcA residues of grass GAX and lignin are also likely to exist, but their presence and importance have not been investigated. Therefore, both feruloylated-Ara*f* and GlcA substitutions are implicated in the cross-linking of wall components and affect wall assembly.

Xylosyl Arabinosyl substitution of Xylan 1 (XAX1), a member of the GT61 glycosyltransferase family, is linked to hydroxycinnamate esterification and to Xyl*p*-Ara*f* substitution of xylan in rice (Chiniquy *et al*., 2012). We recently proposed that XAX1 functions as a glycosyltransferase adding hydroxycinnamic acid-modified Ara*f* substitutions to xylan since xylan from rice *xax1* mutants exhibits a reduction in both hydroxycinnamate-modified Ara*f* monosaccharide and Xyl*p*-Ara*f* disaccharide substitutions (Feijao *et al*., 2022). This hypothesis was supported by the identification of another member of the GT61 family that acts as a xylan Ara*f* 2-*O*-xylosyl-transferase to generate the Xyl*p*-Ara*f* disaccharide substitutions (Zhong *et al*., 2022). Rice *xax1* exhibited changes in cell wall cross-linking, demonstrated by reduction of diverse hydroxycinnamate–hydroxycinnamate and monolignol– hydroxycinnamate coupling products (Feijao *et al*., 2022). Therefore, XAX1 plays a fundamental role in xylan cross-linking but little is known about further consequences for cell wall assembly.

In Arabidopsis, the addition of GlcA substitutions onto the xylan backbone is mediated by Glucuronic acid substitution of xylan (GUX)-1 and GUX2 glycosyltransferases of the GT8 family (Mortimer *et al*., 2010). Arabidopsis *gux1* and *gux2* single mutants have reduced GlcA decorations on different regions of the xylan backbone, whilst the *gux1gux2* double mutant has almost no detectable GlcA substitutions (Mortimer *et al*., 2010; Bromley *et al*., 2013). Furthermore, loss of GlcA in *gux1gux2* xylan results in major improvements in saccharification efficiency, suggesting possible alterations to the molecular architecture of secondary cell walls (Lyczakowski *et al*., 2017). However, it is still not fully understood how GlcA influences interactions between xylan and the lignin and cellulose components of the wall.

To investigate the effect of hydroxycinnamic acid modified-Ara*f* and GlcA substitutions of xylan on wall crosslinking and assembly we chose *Brachypodium distachyon* orthologs of *OsXAX1* (Chiniquy *et al*., 2012) and *AtGUX2* (Mortimer *et al*., 2010) as targets for functional analysis: *BdXAX1A*/Bradi1g06560; *BdXAX1B*/Bradi3g11337 and *BdGUX2*/Bradi4g33330. We used CRISPR/Cas9 mutagenesis to create knockout mutations in *BdXAX1A*, *BdXAX1B* and *Bd*GUX2 and investigated effects on cell wall architecture in these mutants by cryo-SEM, saccharification assays, subcritical water extractions (SWE) and ssNMR. Our data revealed that XAX1A and GUX2 modify distinct types of xylan (Tryfona *et al*., 2023), and demonstrate that loss of XAX1A or GUX2 function induces substantial alterations in the molecular architecture of the wall.

## Materials and Methods

### Phylogeny

All GT61 Clade A and GT8 sequences were downloaded as an orthologous cluster from the comparative genomics platform Plaza Monocots 4.5 (Van Bel et al., 2018), using *Oryza sativa* (rice) GT61 Clade A XAX1 and *Arabidopsis thaliana* GT8 GUX2 protein sequence as queries, respectively. Sequences were aligned with MUSCLE and truncated to their predicted GT domains using a custom Python script (https://www.python.org/). MEGA-X software (Tamura *et al*., 2021) was used to determine an appropriate substitution model (ML), and to build the tree with 100 bootstraps.

### CRISPR constructs and transformation and regeneration of callus

Guide RNAs targeting *BdXAX1a* (Bradi1g06560), *BdXAX1b* (Bradi3g11337), and *BdGUX2*(Bradi1g72350) were identified using the CRISPR-Plant database (genome.arizona.edu/crispr) (Xie et al., 2015). Target sites with low probabilities of mis-targeting were chosen near the 5′ end of the coding sequence. For targeting *BdXAX1a* and *BdXAX1b*, oligonucleotide pairs (pCTA2826/pCTA2827 and pCTA2828/pCTA2829; respectively (Table S1)) were designed to create single gRNAs. In the case of *BdXAX1a*, the double stranded DNA break (DSB) was targeted between nucleotides 7 and 8 of the coding sequence (cds) (Fig. S1c). For *BdXAX1b*, the DSB was targeted between nucleotides 69 and 70 of the cds. For each primer pair, the sequence GGCA was added to the 5’ ends of the forward primer, and AAAC was added to the 5’ end of the reverse primer to create BsaI overhangs for cloning into pRGEB32. For targeting *BdGUX2*, partially complementary and overlapping oligos pCTA2572 and pCTA2573 (Table S1) were designed to create a single gRNA targeting a double-stranded break repair in Bradi1g72350 between nucleotides 40 and 41 of the cds (Fig. S1d). A control gRNA sequence targeting eGFP was designed using pCTA2390 and pCTA2391 (Table S1); this sequence was checked for a low probability of off-target effects against the Bd21-3 genome. Cloning of gRNAs into pRGEB32 was performed as previously described (Petrik, 2020; Petrik *et al*., 2020). Embryogenic callus generation, transformation and regeneration were performed according to previously established methods (Petrik, 2020; Petrik *et al*., 2020).

### Genotyping of XAX1a, XAX1b, and GUX2 CRISPR plants

Genomic DNA was isolated from leaf disks of juvenile CRISPR plants using the Sigma-Aldrich Plant Extract n Amp DNA isolation kit according to manufacturer’s instructions. PCR amplification products using genotyping primers that spanned DSB sites (Table S1) were purified, sequenced and screened for the presence of inserted or deleted nucleotide(s) at these locations.

### Plant growth, growth measurements and ^13^C-labelling of Brachypodium plants

Seeds were stratified in darkness for 48 h at 4 °C, transferred onto soil (Advance M2, ICL Levington) and grown under long day conditions (21 °C, 100 μmol m^-2^ s^-1^, 16 h light/8 h dark). Plant growth was recorded by measuring the length of the tallest tiller for each plant to the tip of the stem.

Rockwool substrate (Grodan, Vital slabs) and an in-house growth medium as described by (Duan *et al*., 2021) were used for hydroponic growth. Stratified seeds were directly placed on hydroponic pots and were grown for 2 weeks under long-day conditions as described above. Accordingly, plants were placed in a sealed [^13^C] enrichment chamber and grown for 7 weeks as previously described (Duan *et al*., 2021).

### Preparation of Alcohol Insoluble Residue (AIR) and Hemicellulose extraction

AIR was prepared as previously described (Tryfona *et al*., 2023). AIR preparations (100 mg) were depectinated in ammonium oxalate (0.5% w v^-1^) at 85 °C for 2 h and xylan was extracted with 4M NaOH according to previous protocols (Tryfona *et al*., 2023).

### Enzymatic hydrolysis and enzymes

Enzymes used in this study were: GH10 endo-β-1,4-xylanase *Cj*GH10 from *Cellvibrio japonicus* (Charnock *et al*., 1998); GH115 α-glucuronidase *Bo*GH115 from *Bacteroides ovatus* (Ryabova *et al*., 2009; Rogowski *et al*., 2014); GH62 α-arabinofuranosidase *Pa*GH62 from *Penicillium aurantiogriseum* (Peng *et al*., 2017), GH3 β-1,4 xylosidase *Tr*GH3 from *Trichoderma reesei* (Margolles-Clark *et al*., 1996) and GH3 xylosidase *Cg*GH3 from *Chaetomium globosum* (NS39127) were gifts from Novozymes (Tryfona *et al*., 2019); endo-glucuronoxylanase *Ec*GH30 from *Erwinia chrysanthemi* (Urbániková *et al*., 2011) and endo arabinoxylanase *Ct*GH5 from *Acetivibrio thermocellus* (formerly *Clostridium thermocellum*) (Correia *et al*., 2011), were gifts from Professor Harry Gilbert (Newcastle University). All enzymes were added at a final concentration of 2 μM and incubated at 21 °C (40 °C for *Ct*GH5) under constant shaking for 24 h.

Five hundred micrograms of extracted xylan were used for enzymatic hydrolysis. Xylan was hydrolysed in 50 mM ammonium acetate buffer pH 6.0 (pH 4.0 for *Ct*GH5) overnight before boiling for 30 min to heat inactivate enzymes. After digestion, samples were taken to dryness *in vacuo* before further processing dependent upon analytical technique used.

### Polysaccharide analysis by carbohydrate gel electrophoresis (PACE)

Derivatization of carbohydrates was performed as previously described (Wilson *et al*., 2022). Carbohydrate electrophoresis and PACE gel scanning and quantification were performed as described (Goubet *et al*., 2002; Goubet *et al*., 2009).

### Mild acid hydrolysis and solid phase extraction (SPE)

Mild acid hydrolysis was performed on 5 mg AIR samples using 50 mM TFA. This procedure was carried out at 100 °C in a heat block for 240 min according to established protocols (Feijao *et al*., 2022). Hydroxycinnamic acid-modified Ara*f* xylan side chains were purified on Sep-Pak C18 Cartridges as previously described (Feijao *et al*., 2022).

### Procainamide derivatisation and Mass spectrometry

Eluted 20% ethanol fractions from SPE purification were resuspended in water and manually spotted on a MALDI plate with procainamide hydrochloride, overlaid with 1 μl 2,5-DHB matrix (10 mg ml^-1^ in 50% aqueous methanol) and analyzed by MALDI-ToF/ToF–MS on an UltrafleXtreme (Bruker) as described previously (Feijao *et al*., 2022).

### Hydroxycinnamate analysis

Hydroxycinnamate analysis was done by saponification of phenolic compounds followed by HPLC analysis (triplicate) as previously described (Rudjito *et al*., 2019). Resulting peaks were compared to external standards of caffeic acid, *p*-coumaric acid, *trans*-ferulic acid, sinapic acid, *trans-*cinnamic acid, and 8,8′ ferulic dehydro-dimer and 5,5′ ferulic acid dehydro-dimer (0.005–0.1 mg ml^-1^).

### Cell-wall sequential fractionation

Cell-wall sequential fractionation was performed as previously described (Tryfona *et al*., 2023).

### Trifluoroacetic Acid Hydrolysis, Sulphuric acid hydrolysis and HPAEC-PAD

For monosaccharide compositional analysis, samples (1 mg) were hydrolysed in 2 M trifluoroacetic acid for 1 h at 120 °C, centrifuged at 15,000 *g* for 5 min and supernatants were reserved for compositional analysis of non-cellulosic polysaccharides, while pellets were reserved for cellulosic glucose analysis. Glucose in cellulose was measured after hydrolysis of TFA pellets in 72% sulphuric acid at room temperature for 30 min and 1 M sulphuric acid for 3 h followed by neutralization with excess barium carbonate. Monosaccharides were separated on a Dionex ICS3000 system equipped with a PA20 column, a PA20 guard column, and a borate trap (Dionex) as previously described (Tryfona *et al*., 2012).

### Lignin quantification

Total lignin content (acid-insoluble and acid-soluble) was measured using a spectroscopic method as previously described (Lu *et al*., 2021).

### Subcritical water extraction (SWE)

Subcritical water extraction was performed by pressurized hot water using a laboratory accelerated solvent extraction Dionex™ ASE™ 350 according to previously established protocols (Ruthes *et al*., 2017) with alterations: extractions were performed using ∼500 mg of Brachypodium AIR at 160 ℃ and pH 7.0 for 15, 30, 60 and 120 minutes; extracts and residues were dialyzed against water with a 3.5 kDa MWCO membrane for 72 h and freeze-dried.

### Size exclusion chromatography (SEC)

Mass distributions of the alkaline and subcritical water xylan extracts were obtained by SEC coupled to differential refractive index (DRI) and UV detectors (SECurity 1260, Polymer Standard Services, Mainz, Germany) with dimethyl sulfoxide (DMSO)with 0.5% w/w LiBr as mobile phase using a pullulan standard calibration, as previously described (Ruthes *et al*., 2017).

### Biomass saccharification

Biomass suspension preparation and saccharification were carried out as described (Lyczakowski *et al*., 2017). D-Glucose and D-Xylose release from biomass was quantified using commercial kits (Megazyme). Sugar concentration for each experiment was standardised with readings obtained from biomass- and enzyme-only controls. All experiments were performed at least in triplicate.

### Cryo-SEM sample preparation and imaging

Fresh last internode and young leaves of 7-week-old Brachypodium plants were prepared for imaging according to a previously established protocol with some alterations: (i) Sample surfaces were washed briefly with 70% ethanol before freezing to remove cuticular waxes (ii) 1 cm-length leaf sections, taken 1 cm from the adjacent node, were used for cryo-preparation. For the measurements of macrofibril width a total between 140-180 macrofibrils were selected at random on all images analysed, quantified and standardized for the platinum layer applied during the coating as previously described (Lyczakowski *et al*., 2019).

### Sold-state NMR experiments

Solid-state NMR (ssNMR) experiments were performed on never-dried *Brachypodium* whole cell samples include one-dimensional (1D) ^13^C NMR spectra, 2D ^13^C-^13^C double-quantum and single-quantum correlation spectra (INADEQUATE), dipolar-doubled 2D ^1-^H-^13^C dipolar chemical-shift (DIPSHIFT) correlation spectra and water-edited ^13^C NMR spectra, measured as described before (White *et al*., 2014; Duan *et al*., 2021). Details about all the NMR experiments used in this study are presented in Table S3 and Methods S1.

## Results

### *Brachypodium distachyon* has two orthologues of rice *XAX1* and one orthologue of Arabidopsis *GUX2*

Based on sequence similarity to rice *OsXAX1* (Chiniquy *et al*., 2012) and Arabidopsis *AtGUX2* (Mortimer *et al*., 2010), we chose *BdXAX1A* (Bradi1g06550), *BdXAX1B* (Bradi3g11337) (Fig. S1a), and *BdGUX2* (Bradi1g72350) (Fig. S1b) as targets for genome editing-mediated knockout. Arabidopsis GUX2 homologue, was targeted in Brachypodium due to its predicted activity to modify xylan backbone with clustered GlcA residues as recently reported in grasses (Tryfona *et al*., 2023). Guide RNA sequences were designed to target Cas9-mediated double-stranded DNA breaks (DSBs) at sites near the 5’ ends of coding sequences (Fig. S2). Three to four independent transgenic lines per mutation were selected for biochemical analysis (Table S2). Soil-grown 4-6 week old *bdxax1a* mutant plants exhibited stunted growth, while *bdxax1b* plants did not exhibit a growth phenotype (Fig. S3a and b). Brachypodium *gux2* mutant plants grew similar to Wild-type (Fig. S3c and d). *BdXAX1A* is expressed in roots, first and last internode and young leaves (Fig. S4a); expression data for *BdXAX1B* are lacking. *BdGUX2* exhibits higher expression in the first and last internode (Fig. S4b).

### Hydroxycinnamic acid-decorated Ara*f* substitutions and cross-linking of xylan via diferulate linkages are reduced in the *bdxax1a* mutant

To investigate the effects of the *bdxax1* mutation on xylan structure, Alcohol Insoluble Residue (AIR) from young leaves of Wild-type and *bdxax1a* mutant lines was depectinated, saponified and subsequently hydrolysed with xylanases from Glycosyl Hydrolase (GH) families GH5 (*Ct*GH5_34; Fig. 1a), GH30 (*Ec*GH30; Fig. 1b) and GH10 (*Cj*GH10; Fig. 1c) (Lombard *et al*., 2013). Hydrolysis of Wild-type, *bdxax1a* and *bdxax1b* saponified xylan yielded characteristic AXe oligosaccharides (Tryfona *et al*., 2023) with predominantly even degrees of polymerisation (DP) (Fig. 1a, DP 6, 8, 10, 12, 14). Therefore, neither lack of *Bd*XAX1A nor *Bd*XAX1B affect the distribution or abundance of Ara*f* residues on AXe. Wild-type and mutant xylan hydrolysed with GH30 glucuronoxylanase, yielded characteristic GAXc short oligosaccharides (Tryfona *et al*., 2023) with backbone DP between 5-7 (Fig. 1b). In contrast, when hydrolysed with GH10 endoxylanase, the xylan structure in *bdxax1a* mutant lines (but not *bdxax1b*) was found to differ from Wild-type (Fig. 1c). GH10 hydrolysis yielded predominantly Xyl, A^3^X, XA^3^X, A^3^A^3^X and D^2,3^XX oligosaccharides (Fig. 1c, lane 1) as previously described in Miscanthus (Tryfona *et al*., 2019). The relative abundance of D^2,3^XX oligosaccharide, which is known to be feruloylated (Himmelsbach *et al*., 1994), was reduced in *bdxax1a* (all three mutant lines; Fig. 1c, lanes 3, 4 and 5) but not in *bdxax1b* plants (Fig. 1c, lanes 6, 7 and 8).

**Figure 1.**
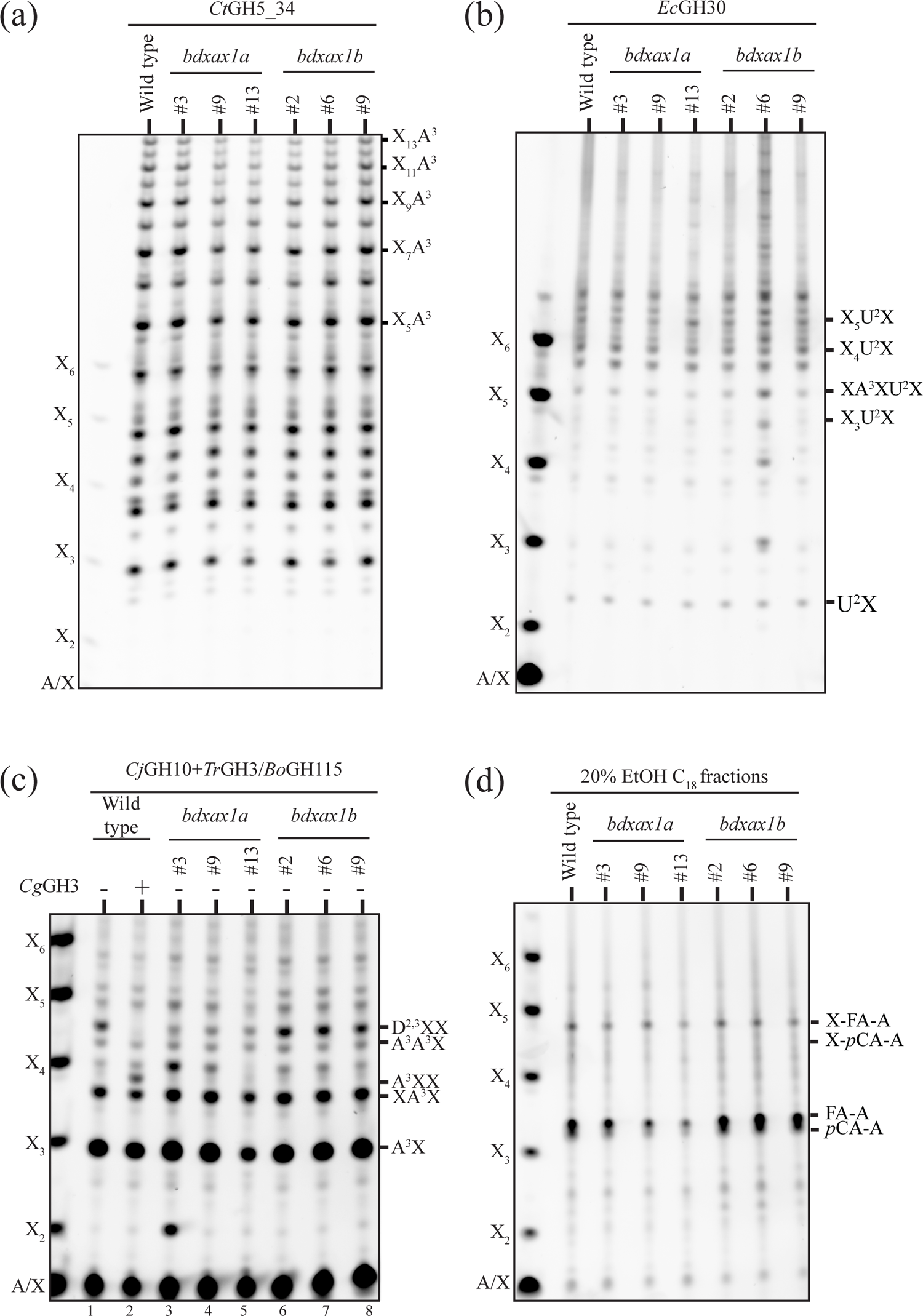
Xylan structure in Brachypodium Wild-type, *bdxax1a* and *bdxax1b* mutant young leaf cell walls. Polysaccharide analysis by carbohydrate gel electrophoresis (PACE) of (a) saponified AIR hydrolysed with GH5 arabinoxylanase; (b) saponified AIR hydrolysed with GH30 glucuronoxylanase; (c) saponified AIR hydrolysed with GH10 endoxylanase and GH115 glucuronidase enzyme cocktail. Confirmation of the assignment of D^2,3^XX oligosaccharide, comes from sensitivity of this structure to a GH3 β-xylosidase (*Cg*GH3) that cleaves terminal Xyl*p* residues from β-Xyl*p*-(1→2)-Ara*f*-(1→3)-Xyl*p* structures (Tryfona *et al*., 2019); (d) mild-TFA hydrolysis products. Hydrolysates were subjected to SPE extraction, and hydroxycinnamic acid modified Ara*f* structures were eluted with 20% EtOH. X, Xyl; A, Ara*f*; U, GlcA; D, Xyl*p*-Ara*f* disaccharide; FA, ferulic acid; *p*CA, *para*-coumaric acid.

To analyse hydroxycinnamic acid modifications of xylan in the Wild-type and mutants we treated AIR with mild TFA to release Ara*f* substitutions. The hydroxycinnamic acid-modified Ara*f* substitutions were separated by solid phase extraction (SPE) using C18 as the stationary phase, elution with ethanol (EtOH) and then analysed by PACE (Fig. 1d) and MALDI-ToF-MS (Fig. S5). PACE analysis yielded three major bands which co-migrated with previously characterised structures: *p*-coumaroyl-Ara*f* (*p*CA-A), feruloyl-Ara*f* (FA-A), and 2-*O*-xylopyranosyl-(5-O-feruloyl)-Ara*f* (X-FA-A) (Feijao *et al*., 2022). 2-*O*-xylopyranosyl-(5-O-*p*-coumaroyl)-Ara*f* (X-*p*CA-A) was also faintly visible. These structures were reduced in *bdxax1a* lines but not in *bdxax1b* (Fig. 1d). Quantification by PACE of Ara*f* monosaccharide and Xyl*p*-Ara*f* disaccharide following alkali release of hydroxycinnamates from these structures revealed a combined reduction of FA-A+*p*CA-A and X-FA-A+X-*p*CA-A in *bdxax1a* (but not in *bdxax1b*) (Fig. 2a). Since *bdxax1b* did not show biochemical phenotypes in the tissues tested we did not continue its characterization. Quantitative analysis by HPLC of mild alkali-extracted hydroxycinnamates revealed significant reduction of *p*CA and FA in all *bdxax1a* lines (Fig. 2b), supporting the finding that Ara*f* hydroxycinnamic acid modifications are affected. In the case of diferulates (diFA), both 5-5’ diFA and 8-8’ diFA were significantly reduced in *bdxax1a*, indicating that cross-linking of xylan chains via diferulate linkages is also affected. This is consistent with findings on rice XAX1 (Feijao *et al*., 2022).

**Figure 2.**
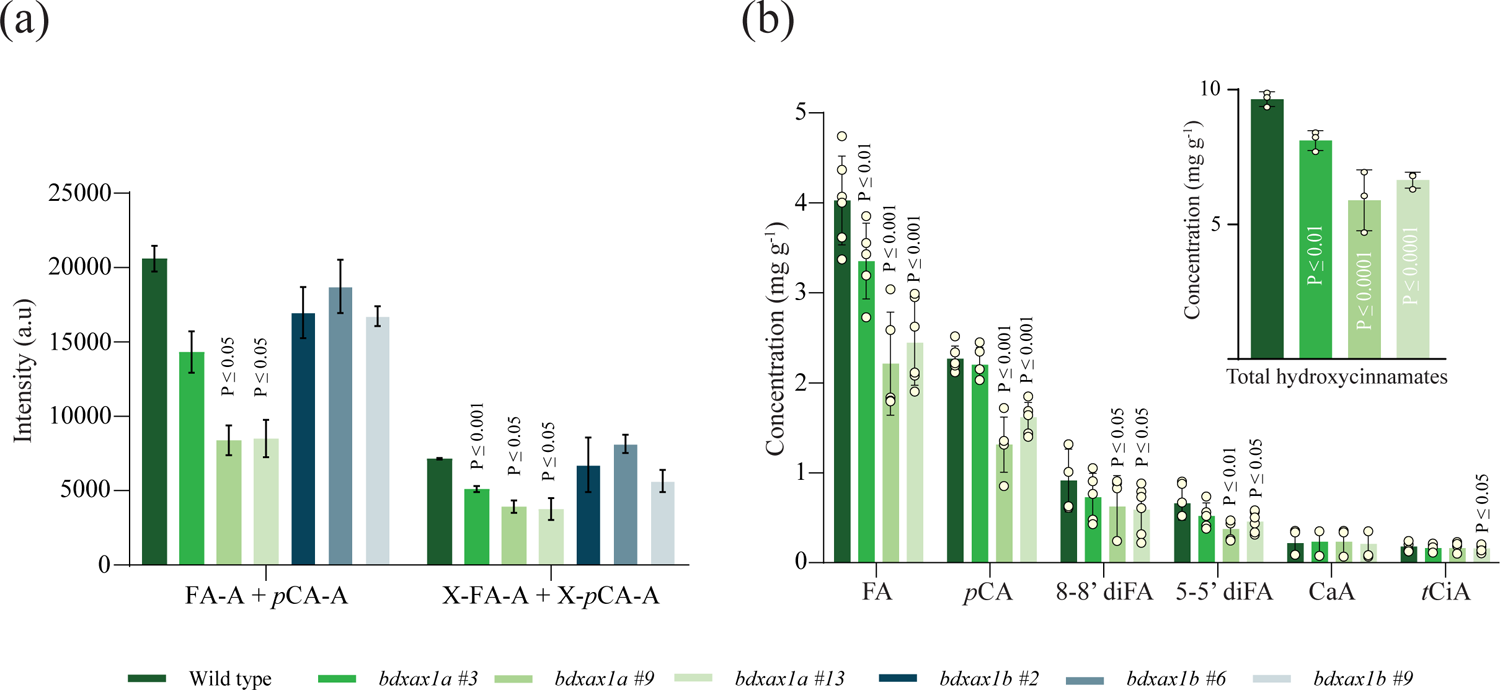
Hydroxycinnamic acid-modified Ara*f* structures are reduced in *bdxax1a*. (a) Quantification of PACE bands (from Fig. 1d) corresponding to FA-A+*p*CA-A and X-FA-A+X-*p*CA-A. Three biological replicates of Wild-type and *bdxax1a* and *bdxax1b* young leaf AIR was TFA-hydrolysed and the SPE C_18_ eluent was analysed by PACE. (b) Quantification of hydroxycinnamates by HPLC. Six biological replicates of Wild-type and *bdxax1a* young leaf AIR were saponified with 2M NaOH, extracted with ethyl acetate, dried, and silylated and analysed by HPLC. X, Xyl; A, Ara*f*; U, GlcA; D, Xyl*p*-Ara*f* disaccharide; FA, ferulic acid; *p*CA, *para*-coumaric acid, CaA, caffeic acid, diFA, diferulic acid; *t*CiA, trans-cinnamic acid. Three replicates of extraction were performed and analysed. Error bars represent standard deviation.

### Reduction of hydroxycinnamic acid-decorated Ara*f* xylan side chains affects the molecular mass and extractability of distinct types of xylan

Since Brachypodium XAX1A facilitates xylan cross-linking, we investigated the interaction of xylan with other wall components by sequentially extracting polysaccharides from Wild-type and *bdxax1a* AIR with a series of chemical treatments (Fig. S6). Interestingly, even though the extractability of AXe and GAXc appeared to be unaffected by loss of *BdXAX1A* function, a proportion of GH10-digestible xylan (presumably hsGAX) was more easily extracted and thus may be less strongly bound to the wall.

Alkali treatment hydrolyses hydroxycinnamates esterified to cell wall polysaccharides (Bach Tuyet Lam *et al*., 1994). Therefore, to identify xylan types more extractable after loss of *Bd*XAX1a while preserving the esterified hydroxycinnamates on Ara*f*, xylan was extracted from Wild-type and *bdxax1a* AIR with subcritical water (SWE) (Ruthes *et al*., 2017; Martínez-Abad *et al*., 2018). Indeed, increased total solids and xylan were extracted from *bdxax1a* compared to Wild-type after 15, 30 and 60 min extractions (Fig. 3a and b, respectively). The extracted xylan in the *bdxax1a* SWE fractions was significantly less substituted with Ara (Ara:Xyl ratio) and had significantly reduced FA and *p*CA content, as well as of 8-8’ and 5-5’ diFA dimers (Fig. 3c, S7a and S7b). Next, we investigated whether *bdxax1a* affected the molecular mass of xylan by means of size-exclusion chromatography (SEC) using simultaneous detection with UV (280 nm) and also refractive index (RI), which non-specifically detects dissolved substances. First, xylan was extracted with 4M NaOH and applied to an SEC column. The molar mass distributions of the xylan did not exhibit substantial differences between Wild-type and *bdxax1a*, indicating that the loss of XAX1A function does not affect the size/degree of polymerization of the xylan polymer (Figure 3d). The relative abundance of the shoulder at smaller molar mass (between 10^3^ and 10^4^ Da; Figure 3d) seems slightly increased in *bdxax1a* as well as in the UV chromatogram (between 15 and 19 ml; Figure S7c), which might correspond to enriched hsGAX population in agreement with our previous study (Tryfona *et al*., 2023). On the other hand, the molar mass distributions of the SW extracts were significantly affected in *bdxax1a* (SWE 30, 60 and 120 min; Fig. 3d). The high molecular mass populations do not exhibit UV absorbance (10-15 ml; Figure S7d), hence can be assigned to starch and/or mixed-linkage glucans (MLGs)). Interestingly, the molecular mass distribution of 30 and 120 min fractions shifts significantly towards lower mass ranges in *bdxax1a* (Figure S7d and f). This could be attributed to a reduction in xylan-xylan cross linking and/or xylan cross linking to lignin, as well as to a higher susceptibility of these fractions to hydrolysis during longer SWE times (Rudjito *et al*., 2019). These data indicate that although *Bd*XAX1A did not affect the length of xylan backbone, it caused significant alterations in xylan cross-linking.

**Figure 3.**
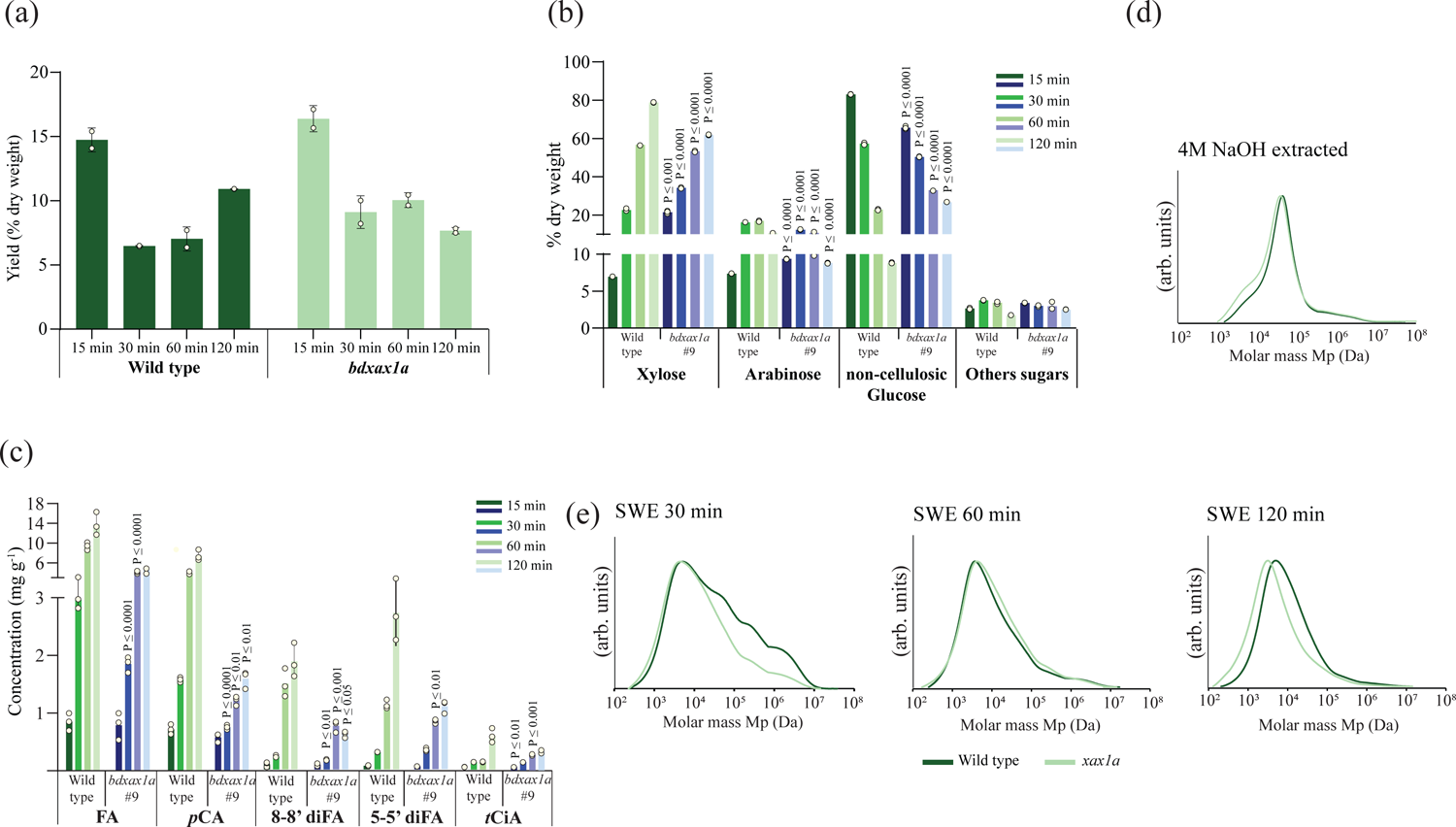
Characterisation of subcritical water extraction fractions (SWE) at different timepoints. Monosaccharide and hydroxycinnamate content were monitored across all fractions, while we assessed the presence of AXe, GAXc and hsGAX in each fraction with xylanases and PACE. (a) Total solid yields (% DW) from Wild-type and *bdxax1a* young leaf samples. Bars represent median and error bars show 95% confidence interval of media for the data; (b) Sugar compositional analysis of SWE fractions for Wild-type and *xax1a* mutant. Three technical replicates for Wild-type and *bdxax1a* young leaf AIR were TFA hydrolysed and analysed by HPAEC-PAD; (c) Quantification of SWE fraction hydroxycinnamates for Wild-type and *bdxax1a* mutant by HPLC. FA, ferulic acid; *p*CA, *para*-coumaric acid, CaA, caffeic acid, diFA, diferulic acid; *t*CiA, trans-cinnamic acid; (d) Molar mass distributions (w log M) of Wild-type and *xax1a* alkali (4M NaOH) extracted xylan and SWE fractions (30 min, 60 min and 120 min) from Brachypodium Wild-type and *bdxax1a* young leaves. Error bars represent standard deviation.

### *Bd*GUX2 modifies a subtype of GAXc

To investigate the role of *BdGUX2* in Brachypodium xylan biosynthesis, AIR was prepared from the last internode of *bdgux2* plants, saponified and characterized with PACE using xylanases GH10 (Fig. 4a) and GH30 (Fig. 4b) in combination with GH62 arabinofuranosidase (*Pa*GH62) to remove Ara*f* side chains and allow easier interpretation of PACE gels. Hydrolysis of Wild-type stem AIR with GH10/GH62 released Xyl and U^2^XX (Fig. 4a). Notably, hydrolysis of *bdgux2* AIR showed a decrease in the intensity of the U^2^XX oligosaccharide for lines 2, 5 and 21, suggesting the mutant contained reduced GlcA substitution of xylan. Quantification of the frequency of GlcA substitution of Xyl by PACE revealed *c.* 35% reduction in *bdgux2* (Fig. 4c), similar to findings for the Arabidopsis *gux2* mutant (Mortimer *et al*., 2010).

**Figure 4.**
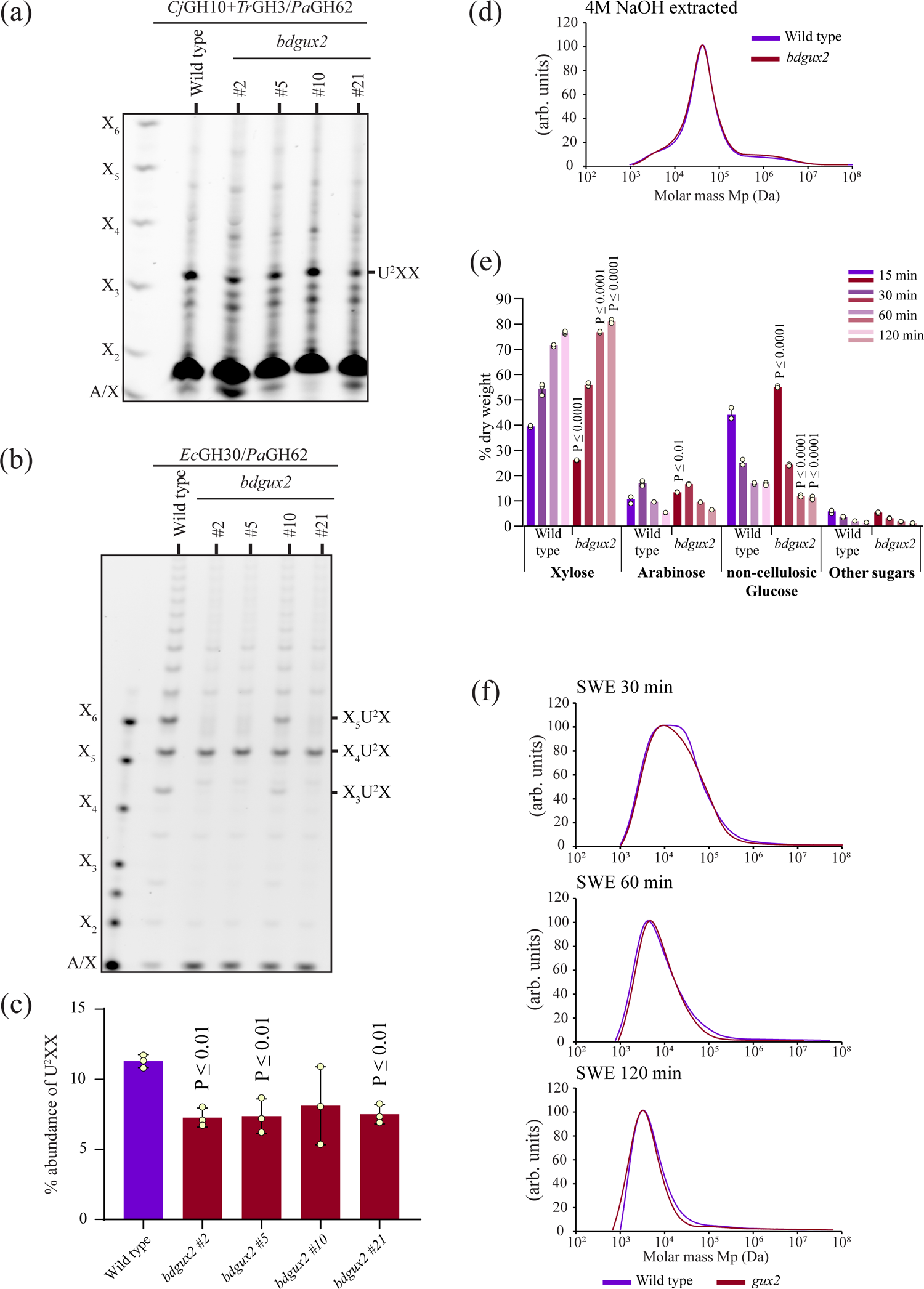
Xylan structure in Brachypodium last internode cell walls. (a) PACE analysis of saponified AIR hydrolysed with GH10 endoxylanase, GH3 xylosidase and GH62 arabinofuranosidase enzyme cocktail; (b) PACE analysis of saponified AIR hydrolysed with GH30 glucuronoxylanase. (c) Quantification of PACE bands corresponding to U^2^XX oligosaccharide. Tree biological replicates of Wild-type and *bdgux2* last internode AIR was saponified, hydrolysed with GH10 endoxylanase, GH3 xylosidase and GH62 arabinofuranosidase enzyme cocktail and analysed by PACE. (d) Molar mass distributions (w log M) of Wild-type and *bdgux2* alkali (4M NaOH) extracted xylan. (e) Sugar composition (% dry weight) for Wild-type and *bdgux2* mutant SWE fractions by HPAEC-PAD. Error bars represent standard error of three Wild-type and mutant technical replicates. (f) Molar mass distributions (w log M) of Wild-type and *bdgux2* SWE fractions (30 min, 60 min and 120 min) from last internodes. X, Xyl; A, Ara*f*; U, GlcA.

To determine any distribution specificity in transfer of GlcA to the xylan backbone by BdGUX2, xylan was hydrolysed with GH30 glucuronoxylanase and GH62 arabinosidase enzyme cocktail. Wild-type xylan yielded characteristic GAXc oligosaccharides (Tryfona *et al*., 2023) with backbone DP between 5 and 7 (Fig. 4b). In contrast, digestion of *bdgux2* xylan released mainly the X_4_U^2^X oligosaccharide and almost no detectable X_3_U^2^X or X_5_U^2^X oligosaccharides. The relative abundance of GlcA oligosaccharides on GAXc estimated by PACE is shown in Table 1. These results indicate that *Bd*GUX2 modifies a subtype of GAXc where GlcA modifications are clustered specifically every 5 or 7 Xyl residues on the xylan backbone, while other unknown enzymes add GlcA spaced every 6 residues to a different region of the xylan backbone.

**Table 1:** Abundance (%) of GlcA modified oligosaccharides on GAXc as revealed by *Ec*GH30 hydrolysis of Brachypodium Wild-type and *bdgux2* mutant line last internode cell walls. Oligosaccharides were quantified from PACE gels using Image J software (n=6); ND, Not Detected.

|  |  | Genotype |  |  |  |
| --- | --- | --- | --- | --- | --- |
|  |  | Wild type | *bdgux2* #2 | *bdgux2* #5 | *bdgux2* #21 |
| Oligosaccharide | X<sub>3</sub>U<sup>2</sup>X | 14.4±1.6 | ND | ND | ND |
|  | X<sub>4</sub>U<sup>2</sup>X | 32.0±1.1 | 72.9±10.6 | 76.1±6.3 | 79.5±6.8 |
|  | X<sub>5</sub>U<sup>2</sup>X | 30.1±1.5 | ND | ND | ND |
|  | X<sub>6</sub>U<sup>2</sup>X | 12.9±0.8 | 17.7±3.2 | 15.9±0.5 | 13.0±4.8 |
|  | X<sub>7</sub>U<sup>2</sup>X | 10.6±0.6 | 9.4±0.8 | 8.0±0.6 | 7.5±0.6 |

### *Bd*GUX2 lack of function alters extractability of distinct domains of xylan

We hypothesised that alteration to GlcA substitution might affect xylan esterification to lignin, which is alkali sensitive. Sequential chemical extractability of AXe and GAXc from AIR appeared unaffected by loss of *Bd*GUX2 function (Fig. S8). The molar mass distribution for the xylan fraction extracted at 4M NaOH did not exhibit any difference between Wild-type and *bdgux2* (Figure 4d and S9d), indicating that *Bd*GUX2 mutation does not affect the length of the xylan backbone.

SWE can isolate native, alkali-sensitive, lignin-carbohydrate-complexes (LCCs) such as lignin esterified to GlcA (Martínez-Abad *et al*., 2018). Therefore, cell wall macromolecules were extracted with SWE and the molecular mass distribution of the fractions (30, 60 and 120 min) was analysed with SEC. The DRI profiles exhibited broad asymmetric peaks for all SW extracts, indicating a dispersity in the molar mass of the isolated xylan populations (Figure S9c-e). More xylan was extracted from *bdgux2* compared to the Wild-type at later extraction times with small differences in molecular mass distribution for the two genotypes (Figure 4e and f). Higher xylan extractability could be a consequence of reduced xylan-lignin cross linking due to decreased GlcA residues on *bdgux2* xylan, or more degradation of material due to higher accessibility of the wall to subcritical water. Interestingly, the *bdgux2* 30 min SWE fraction showed no change in xylan content but exhibited a peak shift towards lower molecular masses compared to the Wild-type extract. Since the longer SW extracts (60 and 120 min) do not exhibit such large differences in molar mass distributions, hydrolytic degradative effects are unlikely. This indicates that lack of *Bd*GUX2 function enables higher extractability of a GAXc population with moderately high molar mass *c.* 100 kDa that is extractable at longer times (60-120 min).

### Reduction of xylan-xylan and xylan-lignin interactions increases the dimensions of Brachypodium macrofibrils

To further investigate the effects of altering xylan crosslinking on the molecular architecture of secondary cell walls, we employed structural imaging of cryo-preserved walls with Scanning Electron Microscopy (cryo-SEM; Fig. 5a-e), previously observed in eudicots and gymnosperms (Lyczakowski *et al*., 2019). Macrofibrils were clearly observed in Brachypodium metaxylem vessels, allowing the measurement of macrofibril diameter. Although there was wide variation in measurements, the mean macrofibril diameter in metaxylem vessels (Fig. 5a and b) was estimated to be 15.8 nm in young leaves and 18.0 nm in last internodes, indicating that Brachypodium macrofibril diameter is comparable to Arabidopsis (Lyczakowski *et al*., 2019). The median macrofibril diameter was found to be *c.* 13% and 16% larger for *bdxax1a* and *bdgux2*, respectively compared to Wild-type (Fig. 5f and g). This indicates that reduction in xylan cross-linking results in wider macrofibrils, a potential indication of a more porous wall or more cell wall material in each macrofibril.

**Figure 5.**
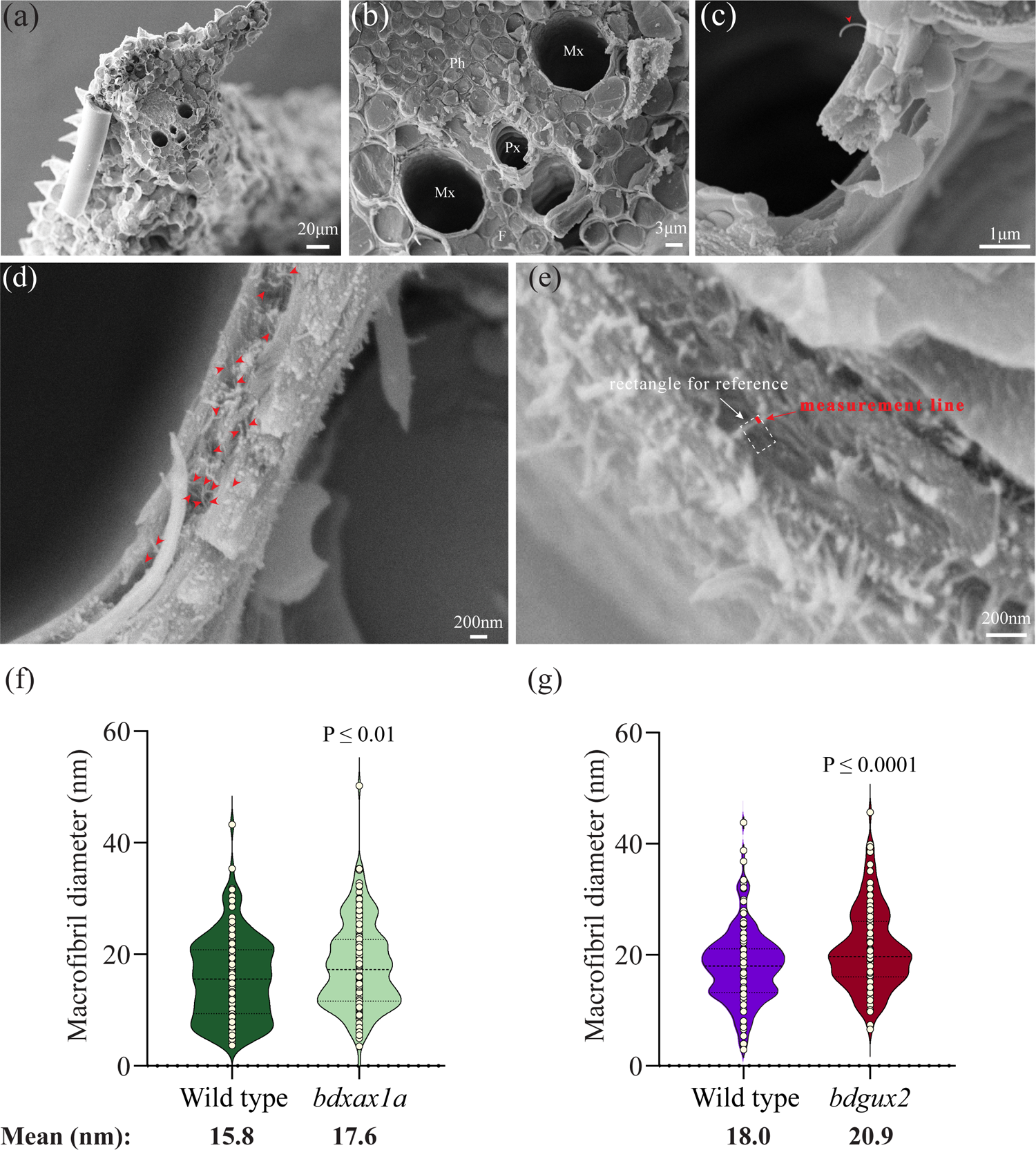
Measurements of Brachypodium cell wall nanofibrils. Cryo-SEM analysis of young leaf sections (a-d) representative images at different magnifications. (b) Tissue anatomy is clearly visible at 10000 magnification. Ph, phloem; Mx, metaxylem; F, fibers; Px, protoxylem. Individual macrofibrils are indicated with red arrows (c and d). (e) Macrofibril diameter was measured at a random site along its length by drawing a rectangle with the long side along the macrofibril and using the perpendicular side as 90° guide to draw a line that span the width of the macrofibril. Scale bars are provided for each image. (f) Cryo-SEM analysis of young leaf sections: Diameter of *bdxax1a* metaxylem cell wall fibrils compared to Wild-type. 140 individual macrofibrils were measured for Wild-type and 174 individual macrofibrils were measured for *bdxax1a* mutant. (g) Cryo-SEM analysis of last internode sections: Diameter of *bdgux2* metaxylem cell wall fibrils compared to wild-type. 167 individual macrofibrils were measured for Wild-type and 140 individual macrofibrils were measured for *bdgux2* mutant. For the measurements at least 3 tissue samples (young leaf of last internode) from each plant and at least 5 plants were used for each genotype.

### Reduced cell wall recalcitrance in *bdxax1a* and *bdgux2*

Cell wall porosity strongly influences the penetration of enzymes into biomass to deconstruct it, and therefore presents an important barrier to hydrolysis (Ding *et al*., 2012). To evaluate the impact of modified xylan branching on grass cell wall assembly, we performed enzymatic saccharification experiments, without any chemical pre-treatment, with biomass from Wild-type and mutant Brachypodium (Fig. 6). There was a significant increase in glucose (Fig. 6a and e) and xylose (Fig. 6b and f) release from both *bdxax1a* and *bdgux2* biomass in comparison to Wild-type. On the other hand, cellulose, xylose/arabinose and total lignin contents were the same (Fig. 4c, d, g and h), indicating that the increased sugar release is not due to increased polysaccharide or reduced lignin content. The improved saccharification suggests *bdxax1a* and *bdgux2* have more porous secondary walls.

**Figure 6.**
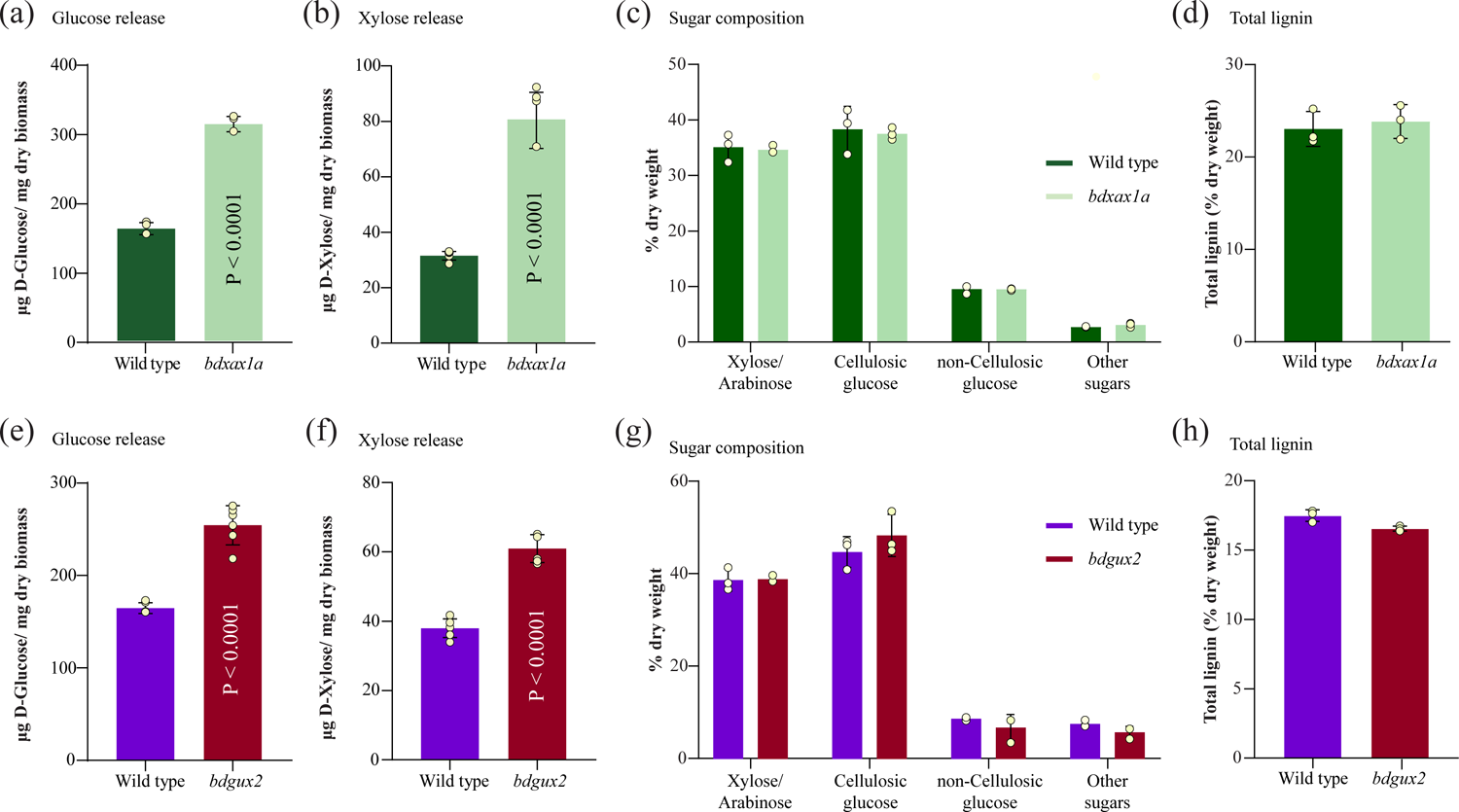
Reduced recalcitrance of biomass lacking hydroxycinnamic acid-modified Ara*f* substitutions. Average (a) D-glucose and (b) D-xylose release following saccharification of Wild-type and *bdxax1a* milled dried biomass. (c) sugar composition (% dry weight) of Brachypodium Wild-type and *bdxax1a* young leaf cell walls (n=3); (d) Total lignin (soluble and insoluble) content is not affected in *bdxax1a* mutant (n=3). Reduced recalcitrance of biomass lacking xylan GlcA-substitutions. Average (e) D-glucose and (f) D-xylose release following saccharification of Wild-type and *bdgux2* milled dried biomass. (g) sugar composition (% dry weight) of Brachypodium Wild-type and *bdgux2* last internode cell walls (n=3); (h) Total lignin (soluble and insoluble) content is not affected in *bdgux2* mutant (n=3). Error bars represent standard deviation of three matching Wild-type and mutant biological replicates of biomass.

### Cell wall molecular architecture is changed in *bdxax1a* and *bdgux2* mutants

To investigate further the architectural changes in the walls of *bdxax1a* and *bdgux2* mutants, we performed ssNMR measurements of never-dried intact Brachypodium tissues. We analysed cell wall composition and architecture using 1D ^13^C ssNMR experiments such as quantitative DP (q-DP), DP with a short recycle delay of 2 s, CP, and INEPT (Fig. S10, S11), and site-resolved 2D ^13^C-^13^C correlation experiments such as 30 ms CORD (CC) (Fig. 7a) and CP-INADEQUATE experiments (Fig. S12). A summary of the different ^13^C chemical shifts is provided in Table S4. These ssNMR spectra show that the overall polysaccharide content is very similar between the Wild-type and mutant. The content of arabinose (Ara) and hydroxycinnamate (FA/*p*CA) species, normalised to internal cellulose, is slightly lower in the *bdxax1a* than in Wild-type, as manifested by the small intensity decrease of the signals at 108.3 ppm (Ara C1) in the quantitative DP, 0.5 ms CP and INEPT spectra (Figure S10a) as well as signals at 169 ppm (FA /*p*CA C9) and 147 ppm (G-lignin) in the q-DP, 0.5 ms CP and 2 s - DP spectra (Fig. S11). This is consistent with the proposed role of *Bd*XAX1 in addition of hydroxycinnamate-modified Ara*f* to the xylan backbone (Feijao *et al*., 2022). Loss of *Bd*XAX1A function also reduced the amount of total and immobilised three-fold xylan (Xn^3f^), as shown by the reduced intensities at 102.4 ppm (Xn^3f^ C1) in the q-DP and 0.5 ms CP spectra (Fig. S10a). The 30 ms CORD provides further support for the decrease of immobilised arabinose and Xn^3f^, manifested by moderate intensity reduction of the diagonal Ara (C1, C1) cross peak at (108.3, 108.3) ppm and the Xn^3f^ (C1, C2/3) cross peak at (102.5, 73.8) ppm (Fig. 7a). On the other hand, the intensity of Xn^2f^ (C4, C5) cross peak at (82.2, 64.3) ppm in 30 ms CORD spectrum remains the same in *bdxax1a* and Wild-type samples (Fig. 7a, 82.2 ppm cross-section). Furthermore, the CP-INADEQUATE spectra (Fig. S12) reveal no significant changes in the content of highly rigid portions of Xn^3f^ and Xn^2f^ upon *BdXAX1a* mutation. The computed ratio between the integrated areas of Xn^3f^ C4 cross-peak at (140.5, 77.3) ppm in *bdxax1a* and Wild-type spectra as well as the ratio between the integrated areas of Xn^2f^ C4 cross-peak at (146.4, 82.2) ppm in these spectra are the same within the experimental uncertainty. Similarly, comparison of Wild-type and *bdgux2* q-DP spectra indicate that loss of *Bd*GUX2 function resulted in a slightly reduced amount of arabinose (Ara C1; 108.3 ppm) compared to Wild-type cell wall (Fig. S10b).

**Figure 7.**
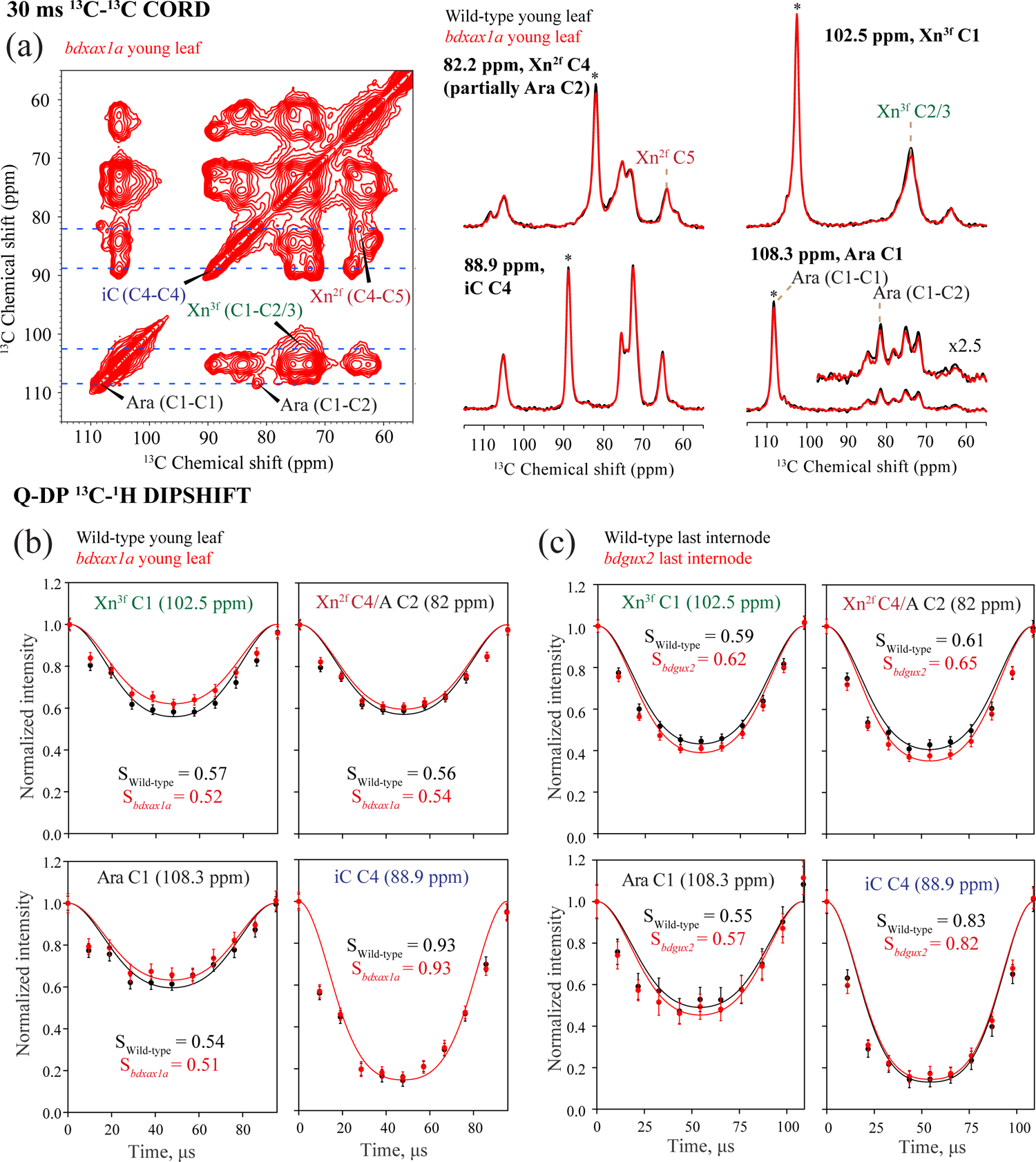
2D ^13^C−^13^C 30 ms CORD and ^13^C^-1^H dipolar-doubled Q-DP DIPSHIFT spectra show changes in polysaccharide content and dynamics. (a) Polysaccharide region of 2D 30 ms CORD of *bdxax1a* young leaf sample, with key cross-sections extracted at the indicated ω_1_ chemical shifts of the Wild-type (black) and *bdxax1a* mutant (red) young leaf cell walls shown on the right. The spectra of the two samples are scaled to match the absolute intensities at 88.9 ppm cross-sections, and the same scaling applied to all the cross-sections shown. Asterisks indicate the diagonal peaks in each cross-section. The inset with 2.5x intensity magnification at the 108.3 ppm cross-section highlights the Ara (C1-C2) signal. The polysaccharide region of 2D 30 ms CORD of Wild-type young leaf sample is not shown due to similarity to the *bdxax1a* spectra. (b) Quantitative DIPSHIFT ^13^C−^1^H dipolar dephasing curves for selected ^13^C sites in Wild-type (black) and xax1a (red) young leaf cell walls. The H-C dipolar order parameters S_sample_ from best-fit simulations are indicated in each panel. The Xn^3f^ xylan C1 peak at 102.5 ppm shows weaker dipolar coupling for the *bdxax1a* mutant than the Wild-type cell wall, while other polysaccharides do not exhibit significant mobility differences between the two samples. The observed differences in the dephasing curves of Wild-type and xax1a of Ara C1 at 108.3 ppm are within noise level. (c) Quantitative DIPSHIFT ^13^C−^1^H dipolar dephasing curves for selected ^13^C sites in the last internode Wild-type (black) and *bdxax1a* (red) cell walls. The best-fit C-H dipolar order parameters S_sample_ are indicated in each panel. The Xn^3f^ xylan C1 peak at 102.5 ppm as well as the Xn^2f^ C4 and arabinose C2 mixed peak at 81.6 ppm show weaker dipolar coupling for the *bdgux2* mutant than the Wild-type cell wall, while the other polysaccharides do not exhibit significant mobility differences. Error bars in panels (b) and (c) are the uncertainties propagated from the spectral noise. Representative cross sections of these spectra are shown in Fig. S14.

To investigate further the impact of *bdxax1a* or *bdgux2* mutations on the dynamics of wall polysaccharides, we conducted quantitative and CP-based ^13^C-^1^H DIPSHIFT experiments (Fig. 7b-c, Fig. S13 and Fig. S14). Analysis of the resulting dipolar dephasing curves revealed a decrease in the C-H dipolar order parameter for both total (Fig. 7b, 102.5 ppm) and immobilised Xn^3f^ (C1, 102.5 ppm, Fig. S13a) in the *bdxax1a* mutant compared to Wild-type, consistent with an increase in mobility. However, no significant change in the dynamics of arabinose (Ara C1 108 ppm, Fig. 7b and Fig. S13a) was observed for the *bdxax1a* mutant. In contrast, we found that in the *bdgux2* mutant the C-H dipolar order parameters for both total and immobilised Xn^3f^ (C1, 102.5 ppm; Fig. 7c and Fig. S13b) and immobilised arabinose (Ara C1, 108 ppm; Fig. S13b) have increased compared to the Wild-type, indicating a decrease of mobility. Neither *bdxax1a* nor *bdgux2* mutations have significant impact on the dynamics of cellulose (iC C4; 89 ppm) or Xn^2f^ (82 ppm).

To determine if *bdxax1a* and *bdgux2* mutations induce any changes in water accessibility of cell wall components, we performed a water-edited CP experiment (Fig. 8). In this experiment, the signal coming from the magnetization of water protons diffuses to nearby carbons so that at short diffusion times, the signal from carbons closest to water builds up first (Cresswell *et al*., 2021). Interestingly, the resulting spin diffusion intensity build-up curves for young leaf (Fig. 8b) and last internode (Fig. 8d) samples show faster intensity build-up for the mutants than the Wild-type, indicating increased water accessibility of both *bdxax1a* and *bdgux2* cell walls compared to their respective Wild-type tissues. This finding is consistent with an increase in the porosity of the cell wall in both mutants.

**Figure 8.**
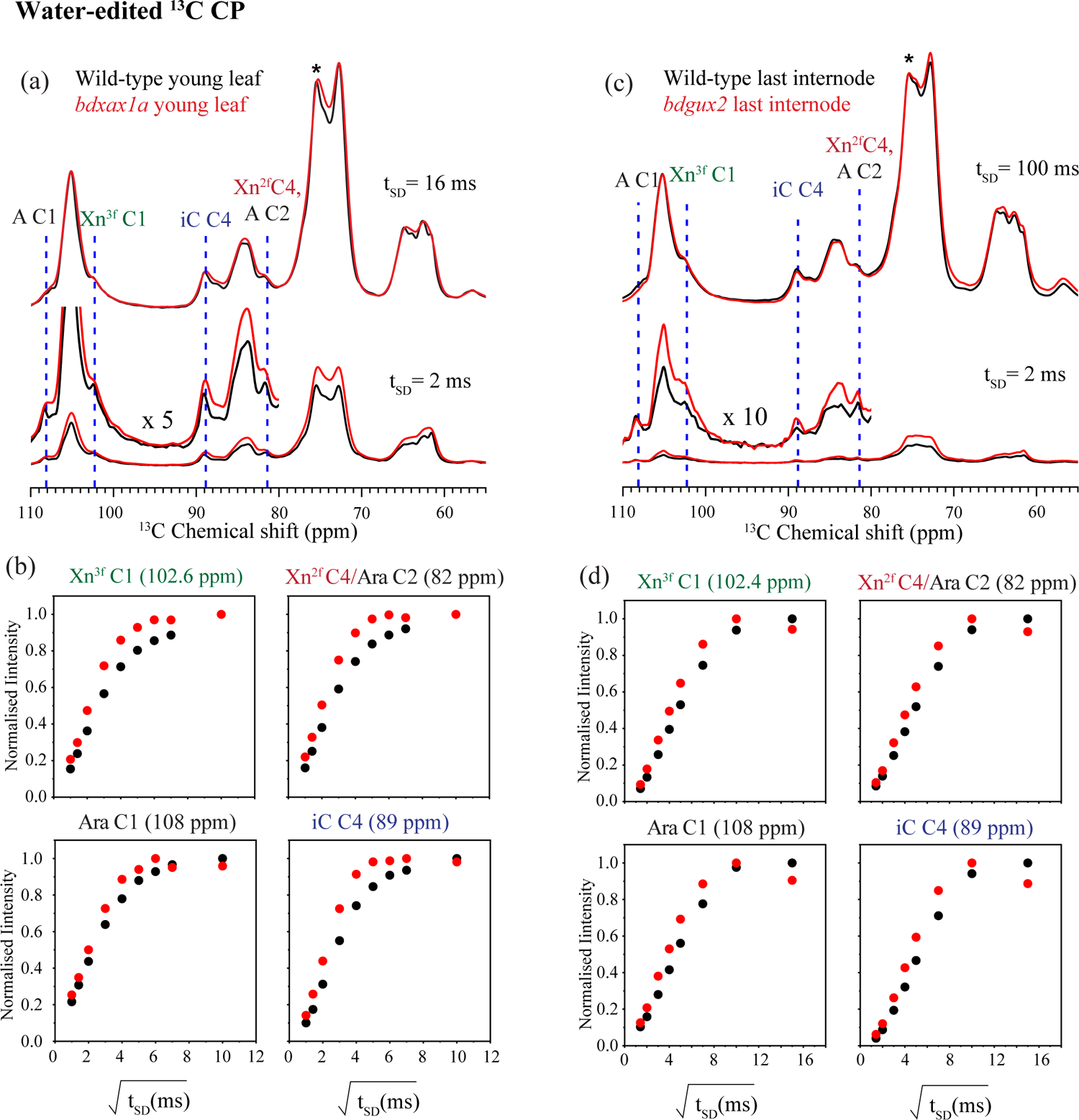
Water-accessibility of Brachypodium secondary cell wall. (a) Comparison of the 1D water-edited ^13^C NMR spectra of never-dried Brachypodium Wild-type (black) and *xax1a* (red) young leaves for 2 ms and 16 ms ^1^H spin diffusion mixing times. Mutant spectra were simultaneously scaled to match the signal of interior cellulose iC C4 at 89 ppm (indicated with an asterisk (*)) in long (16 ms) diffusion time spectrum of Wild-type. Assignments of the major cell wall components are indicated for arabinose (Ara) carbon 1 (C1) and C2, three-fold xylan (Xn^3f^) C1, interior cellulose (iC) C4; two-fold xylan (Xn^2f^) C4. The *bdxax1a* mutant sample has higher signal intensity than Wild-type at short (2 ms) diffusion time. (b) Representative water-to-polysaccharide ^1^H transfer build up curves for Brachypodium young leaf and *xax1a* mutant. The data for arabinose (Ara) carbon 1 (C1), carbon 2 (C2), three-fold xylan (Xn^3f^) C1, interior cellulose (iC) carbon 4 (C4), surface cellulose (sC) C4, and two-fold xylan (Xn^2f^) C4 are compared between Wild-type and *bdxax1a* mutant. All mutant polysaccharides have faster build-up due to their enhanced contact with water. (c) Comparison of the 1D water-edited ^13^C NMR spectra of never-dried Brachypodium Wild-type (black) and *gux2* (red) last internodes for two spin diffusion mixing times, 2 ms and 100 ms. The *bdgux2* mutant sample has higher signal intensity than Wild-type at short mixing time. (d) Representative water-to-polysaccharide [^1^H] transfer build up curves for Brachypodium last internode and *gux2* mutant. The data for arabinose (Ara) C1, C2, three-fold xylan (Xn^3f^) C1, interior cellulose (iC) C4, surface cellulose (sC) C4 and two-fold xylan (Xn^2f^) C4 are compared between Wild-type and *bdgux2* mutant. All mutant polysaccharides have faster build-up due to their enhanced contact with water.

## Discussion

Xylan in grass cell walls was recently demonstrated to consist of distinct types with different substitution patterns (Tryfona *et al*., 2023). We hypothesised that these xylan populations reflect functional specialisation with different interactions with cellulose and lignin. By altering the structure of specific xylan domains using genetic tools we sought to gain useful insights into their functions. In this work we specifically targeted Brachypodium *Bd*XAX1 and *Bd*GUX2 enzymes that likely add hydroxycinnamate ester modified Ara*f* residues (Feijao *et al*., 2022) or GlcA decorations (Mortimer *et al*., 2010), respectively, to xylan. The mutants show changes in cell wall architecture and assembly that lead to increased accessibility of the polymers to water and hydrolytic enzymes.

*Bd*XAX1A is required for some of the hydroxycinnamic acid modifications on Ara*f* xylan side-chains in Brachypodium young leaves, with *bdxax1a* exhibiting significantly reduced hydroxycinnamic acid content including diFAs, consistent with our recent work on the rice *xax1* mutant (Feijao *et al*., 2022). Interestingly, sequential cell wall extraction and SWE demonstrated that a specific type of xylan was more extractable in the *bdxax1a* mutant. We postulate that during biosynthesis *Bd*XAX1A modifies the GH10-digestible primary cell wall hsGAX, since in the mutant we observed a significant reduction of the D^2,3^ xylan structure which is more abundant in grass leaves (Tryfona *et al*., 2019). *Bd*XAX1A might also be involved in biosynthesis of another xylan type, AXe. Most importantly, xylan in *bdxax1a* SWE fractions exhibited a significant reduction in molecular mass distribution compared to Wild-type even though the molecular mass distributions of alkali-extracted xylans appeared to be very similar. Therefore, loss of *Bd*XAX1A function likely reduces the mass of xylan-xylan and/or xylan-lignin complexes. During wall maturation, cross-linking of xylan chains occurs through ferulate oxidative coupling mediated by peroxidases and laccases (Buanafina & Morris, 2022). We propose that as cells mature, ferulate from hsGAX and/or AXe crosslinks with other xylan chains via diFA bridges or with lignin via ferulate-mediated LCCs, resulting in cessation of elongation, initiation of lignification and fortification of the wall. Indeed, wall-bound peroxidases coincide with the formation of diFAs and cessation of elongation growth in maize coleoptiles (Wakabayashi *et al*., 2012). Furthermore, xylan cross-linking has been reported to occur in the S2 layer of bamboo SCWs (Munekata *et al*., 2022). An alteration of primary cell wall hsGAX crosslinking could explain the dwarfed phenotype of *bdxax1a* plants, as these walls may not be able to modulate sufficiently their mechanical properties (Bidhendi & Geitmann, 2016) due to reduced FA-mediated cross-linking in the wall affecting the process of cell growth.

*Bd*GUX2 modifies a subtype of GAXc where GlcA modifications are clustered on the xylan backbone, whilst another unknown GUX enzyme (perhaps more closely related to Arabidopsis GUX1 and GUX3) is responsible for evenly distributed GlcA residues. This discovery confirms and extends the findings of different types of xylan in grass walls (Tryfona *et al*., 2023) because it revealed that GAXc has domains that have clustered GlcA six residues apart, distinct from the *Bd*GUX2 modified xylan. Loss of *Bd*GUX2 function does not have an effect on plant growth, similar to a lack of growth defects in Arabidopsis *gux1gux2* plants (Mortimer *et al*., 2010). The existence of GlcA-mediated xylan-lignin cross linking in grasses, and the importance for wall assembly, was previously unexamined. Consistent with their presence, we found that the molecular mass distribution of SWE fractions was reduced in *bdgux2,* even though the molecular mass distribution of alkali extracted xylan from Wild-type and *bdgux2* appeared the same. These data implicate GlcA substitutions on grass xylan in LCC formation.

We considered that decreased crosslinking of xylans might alter cell wall architecture. Wild-type Brachypodium macrofibril diameters were broadly similar to those reported for maize using Atomic Force Microscopy (AFM) (Ding *et al*., 2012) and also to those reported for eudicots using cryo-SEM (Lyczakowski *et al*., 2019). The increase in fibril diameter in *bdxax1a* and *bdgux2* over the Wild-type may arise from the reduction of xylan crosslinking and hence increased porosity of the wall. Further evidence of increased wall porosity arose from saccharification assays. A key factor that impacts wall degradation is the access of enzymes to their substrates, as exemplified by enzymatically modified kraft pulp fibers (Mansfield *et al*., 1997; Esteghlalian *et al*., 2001). We found enhanced saccharification in *bdxax1a*, caused presumably by the significant reduction of diferulate bridges and Ara*f*-lignin links that maintain a compact structure that interferes with access of hydrolytic enzymes. Loss of *Bd*GUX2 function also resulted in increased saccharification, supporting the notion that xylan and lignin in the cell wall is crosslinked via GlcA esters.

Consistent with the proposed increased cell wall porosity in *bdxax1a* and *bdgux2* mutant tissues, our ssNMR water-edited ^13^C CP data demonstrate that the walls of *bdxax1a* and *bdgux2* mutants are more accessible to water compared to Wild-type. Indeed, loosening of the wall network due to reduced cross-linking is expected to enhance the water accessibility of cell wall biopolymers. Intriguingly, even though both mutations result in increased cell wall porosity, they induce opposite changes in biopolymer dynamics, as revealed by the ^13^C-^1^H DIPSHIFT experiments confirming the presence of different types of xylan in grass walls with different functionalities (Tryfona *et al*., 2023). We speculate that FA from hsGAX (and/or AXe) crosslinks with other xylan chains via diFA bridges or with lignin via ferulate-mediated LCCs, whereas GlcA esters facilitate cross-linking of GUX2 modified domain of GAXc with lignin. According to the grass secondary CW model (Tryfona *et al*., 2023), hsGAX is found in the matrix in a three-fold conformation and is relatively rigid due to cross-linking with other xylans and lignin. In this scenario, *bdxax1a* mutation could result in a more dynamic three-fold xylan, due to reduced restrictions in xylan mobility. Extending the proposed grass SCW model (Tryfona *et al*., 2023) we propose that the GUX2 modified domain of GAXc is found in the matrix, while another GAXc domain with GlcA residues evenly spaced every 6 xyloses is found in the two- or distorted two-fold conformation that is bound to cellulose. Thus, loss of GlcA substitutions on the GUX2 domain would result in reduced cross linking with lignin. In the *bdgux2* mutant, this GUX2 modified GAXc domain may have longer unsubstituted regions that could begin to associate with surfaces of cellulose. This could result in Xn^3f^ becoming more rigid.

In summary, our data demonstrate that both *bdxax1a* and *bdgux2* walls are more porous. Distinct types of xylan crosslink with each other or lignin via distinct xylan substitutions, enabling commelinid monocots to maintain a compact wall structure. Grasses provide the major cereal crops and most of the grazing for domesticated animals and is therefore the most important plant family for human society. Our results demonstrate that removal of FA-Ara*f* and removal of GlcA substitutions can be effective strategies to optimise grass biomass properties for use in a sustainable bioeconomy.

## Acknowledgements

This work was supported as part of The Center for Lignocellulose Structure and Formation, an Energy Frontier Research Center funded by the US Department of Energy, Office of Science, Basic Energy Sciences, under Award number DE-SC0001090. FV acknowledges the financial support from the Swedish Research Council (Project Grant 2020-04720) and the Wallenberg Wood Science Centre through the Knut and Alice Wallenberg Foundation (KAW). DRM is supported by a Margarita Salas grant funded by the European Union NextGenerationEU (RD 289/2021). We thank Dr. Pu Duan for insightful comments on NMR data interpretation.

## Author contributions

T.T. planned and designed the research, performed experiments and analysed all data; Y.P performed ssNMR experiments; Y.P and M.H analysed ssNMR data and contributed to manuscript writing; D.P. generated CRISPR/Cas9 mutants; R.W. acquired Cryo-SEM images; D.R.M performed hydroxycinnamate quantification, SWE and SEC experiments and analysed data with support from F.V.; A.E.P provided help with ^13^C labelled plant growth; X.Y. provided technical support to T.T.; P.K.D performed lignin quantification and H_2_SO_4_ hydrolyses; C.A, T.T. and P.D. conceived original research plans; T.T. wrote the manuscript with support from P.D.; all authors read and commented on the article; P.D. agrees to serve as the author responsible for contact and ensures communication.

## Data availability

The data that support the findings of this study are available from the corresponding author upon reasonable request.

