## Supplemental material for "Altering the substitution and crosslinking of glucuronoarabinoxylans affects cell wall porosity and assembly in *Brachypodium distachyon*"

Article acceptance date: n/a

The following supporting information is available for this article:

**Fig. S1** Phylogenetic analysis.

**Fig. S2** Gene Maps of *Bd*XAX1a, *Bd*XAX1b, *Bd*GUX2 showing location of double stranded breaks (DSBs) induced by Cas9, followed by mutations created by imprecise NHEJ.

**Fig. S3** Plant growth measurements. The growth of Brachypodium mutant plants grown under greenhouse conditions was measured every week and was compared to Wild-type.

**Fig. S4** Gene expression profiles at different Brachypodium developmental stages.

**Fig. S5** MALDI-MS analysis of NaOH-sensitive xylan hydrolysis products of 20% ethanol fraction of the SPE $C_{18}$ eluate for Wild-type and *bdxax1a* mutant lines.

**Fig. S6** Sequential cell wall extraction from Brachypodium Wild-type and *bdxax1a* young leaves.

**Fig. S7** Characterisation of subcritical water extraction fractions (SWE) from mature leaves of Brachypodium Wild type and *bdxax1a* mutant at different timepoints**.**

**Fig. S8** Sequential cell wall extraction from Brachypodium Wild-type and *bdgux2* young leaves.

**Fig. S9** Characterisation of subcritical water extraction fractions (SWE) from last internodes of Brachypodium Wild type and *bdgux2* mutant at different timepoints**.**

**Fig. S10** Comparison of carbohydrate regions of 1D [$^{13}C$] NMR spectra among *Brachypodium* samples.

**Fig. S11** Full 1D $^{13}C$ NMR spectra of *Brachypodium* young leaf samples highlighting lignin region.

**Fig. S12** 2D $^{13}C$–$^{13}C$ CP-INADEQUATE spectra of *Brachypodium* young leaf cell walls.

**Figure S13.** 2D $^{13}C$–$^{1}H$ dipolar-doubled CP-DIPSHIFT spectra of *Brachypodium* young leaf and last internode cell walls.

**Figure S14.** Representative cross sections of the 2D $^{13}C$–$^{1}H$ dipolar-doubled Q-DP DIPSHIFT spectra of *Brachypodium* young leaf and last internode cell walls.

**Table S1.** Summary of genotypes of CRISPR/Cas9 lines.

**Table S2.** List of Primers used in this study.

**Table S3.** Experimental parameters of the $^{13}C$ MAS NMR spectra for Brachypodium samples.

**Table S4.** $^{13}C$ Chemical shift of the major polysaccharide and lignin species.

**Methods S1.** Solid-state NMR experiments.

### Supplementary Figures

**Fig. S1 Phylogenetic analysis** of (a) Rice (*Oryza sativa*) and Brachypodium (*Brachypodium distachyon*) clade A members of the GT61 family. (b) Arabidopsis (*Arabidopsis thaliana*), rice (*Oryza sativa*), maize (*Zea mays*) and Brachypodium (*Brachypodium distachyon*) GT8 family members. The GT61 and GT8 members are shown with their locus identifiers and names if available. *Bd*XAX1A, *Bd*XAX1B and *Bd*GUX2 characterized in this article are highlighted in red.

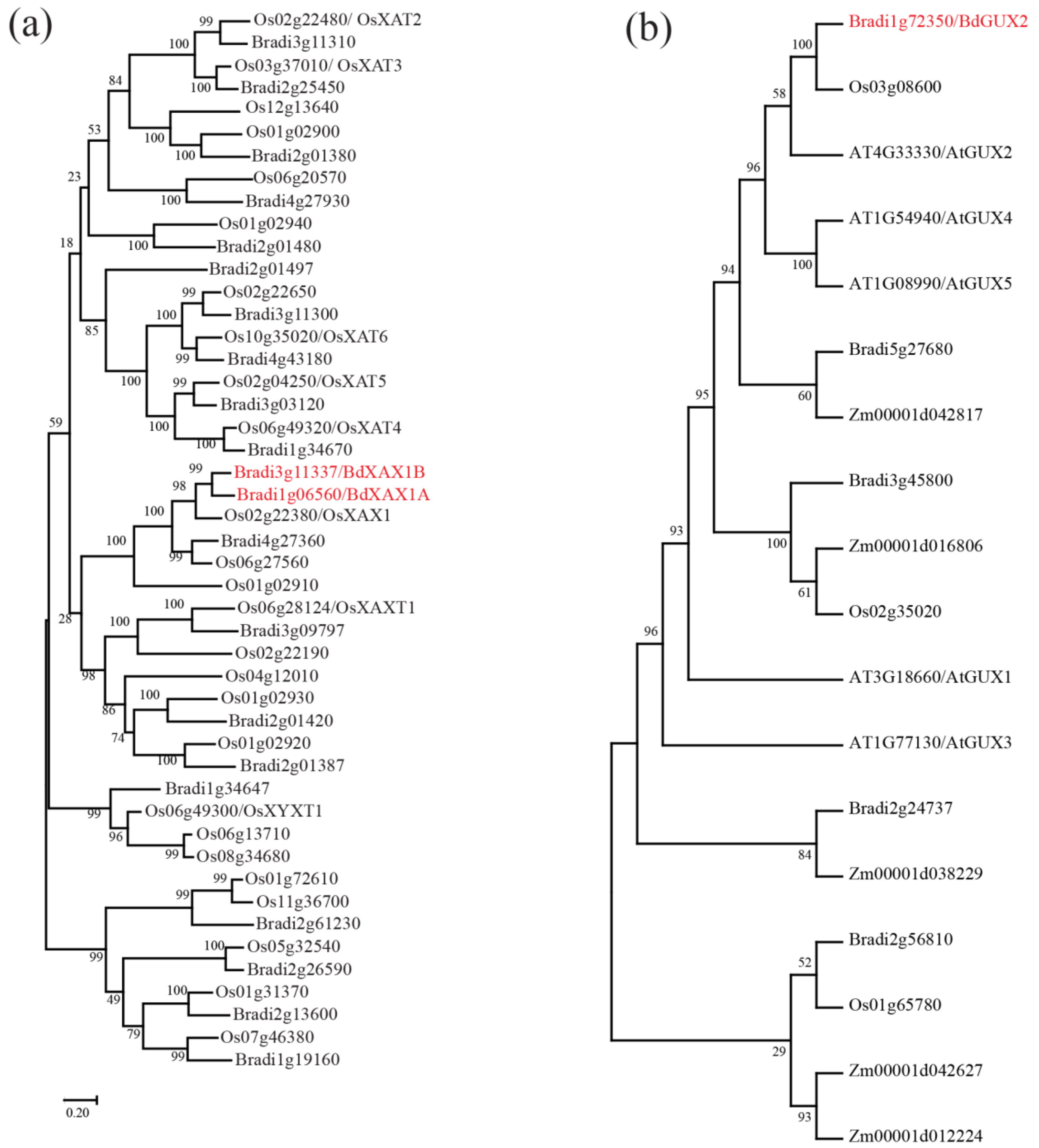
(a)
Os02g22480/ OsXAT2
Bradi3g11310
Os03g37010/ OsXAT3
Bradi2g25450
Os12g13640
Os01g02900
Bradi2g01380
Os06g20570
Bradi4g27930
Os01g02940
Bradi2g01480
Bradi2g01497
Os02g22650
Bradi3g11300
Os10g35020/OsXAT6
Bradi4g43180
Os02g04250/OsXAT5
Bradi3g03120
Os06g49320/OsXAT4
Bradi1g34670
Bradi3g11337/BdXAX1B
Bradi1g06560/BdXAX1A
Os02g22380/OsXAX1
Bradi4g27360
Os06g27560
Os01g02910
Os06g28124/OsXAXT1
Bradi3g09797
Os02g22190
Os04g12010
Os01g02930
Bradi2g01420
Os01g02920
Bradi2g01387
Bradi1g34647
Os06g49300/OsXYXT1
Os06g13710
Os08g34680
Os01g72610
Os11g36700
Bradi2g61230
Os05g32540
Bradi2g26590
Os01g31370
Bradi2g13600
Os07g46380
Bradi1g19160
0.20
(b)
Bradi1g72350/BdGUX2
Os03g08600
AT4G33330/AtGUX2
AT1G54940/AtGUX4
AT1G08990/AtGUX5
Bradi5g27680
Zm00001d042817
Bradi3g45800
Zm00001d016806
Os02g35020
AT3G18660/AtGUX1
AT1G77130/AtGUX3
Bradi2g24737
Zm00001d038229
Bradi2g56810
Os01g65780
Zm00001d042627
Zm00001d012224

FIGURE S1

**Fig. S2 Gene Maps of XAX1a, XAX1b, GUX2 showing location of double stranded breaks (DSBs) induced by Cas9, followed by mutations created by imprecise NHEJ.** (a) Genomic DNA map of XAX1a Bradi1g06560. The total genomic DNA sequence is marked as 2768 nucleotides in length, and the features were marked based on their annotation in Phytozome 13 within the genomic sequence in Bd21-3 genome that is homologous to the Bradi1g06560 gene in the Bd 3.1 reference genome. The double stranded break (DSB) site performed by Cas9 is known to be located between the 3$^{rd}$ and 4$^{th}$ nucleotides 5' of the protospacer adjacent motif (PAM) immediately 3' of the designed guide RNA sequences. The DSB was targeted to occur between the seventh and eighth nucleotides of exon 1. This will induce INDELS to occur within the S3 codon of the cds. Insertions of deletions of 1 or 2 nucleotides will shift the translational reading frame, creating a downstream premature translational stop codon, and thus a truncated and non functional enzyme. (b) Genomic DNA map of XAX1g Bradi1g11337. The total genomic DNA sequence is marked as 4382 nucleotides in length, and the features were marked based on their annotation in Phytozome 13 within the genomic sequence in Bd21-3 genome that is homologous to the Bradi3g11337 gene in the Bd 3.1 reference genome. The double stranded break (DSB) site performed by Cas9 is known to be located between the 3$^{rd}$ and 4$^{th}$ nucleotides 5' of the protospacer adjacent motif (PAM) immediately 3' of the designed guide RNA sequences. The DSB was targeted to occur between the 69$^{th}$ and 70$^{th}$ nucleotides of exon 1. This will cause upon incorrect repair by NHEJ, INDELS to occur between P33 and P34 codons encoding of the amino acid. If one or 2 nucleotides are inserted or deleted at this position, it will shift the translational reading frame, creating a downstream premature translational stop codon, and thus a truncated and non-functional enzyme. (c) Genomic DNA map of GUX2 Bradi1g72350. The total genomic NA sequence is marked as 5825 nucleotides in length, and the features were marked based on their annotation in Phytozome 13 within the genomic sequence in Bd21-3 genome that is homologous to the Bradi1g72350 gene in the Bd 3.1 reference genome. The double stranded break (DSB) site performed by Cas9 is known to be located between the 3rd and 4th nucleotides 5' of the protospacer adjacent motif (PAM) immediately 3' of the designed guide RNA sequences. The DSB was targeted to occur between the 40th and 41st nucleotides of exon 1. This will cause upon incorrect repair by NHEJ to occur within the S14 codon. Indels of one or two nucleotides will shift the translational reading frame, creating a downstream premature translational stop codon, and thus a truncated and non-functional enzyme.

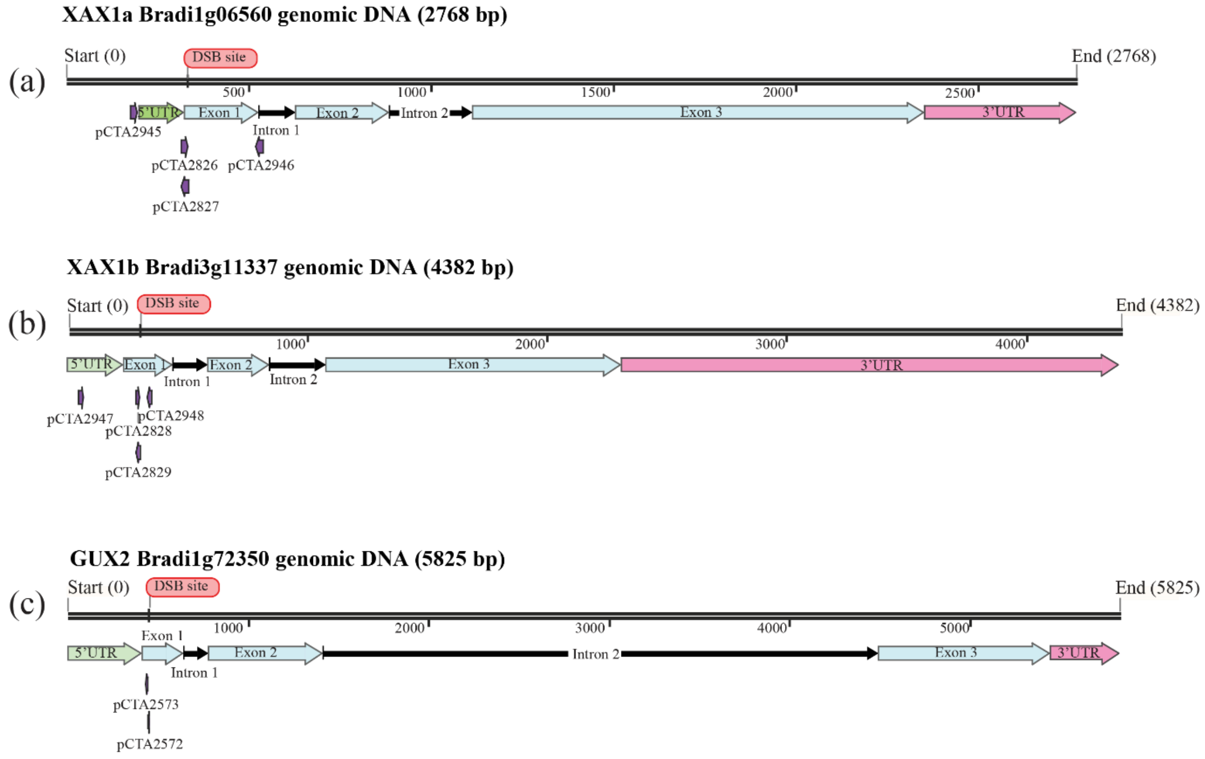


FIGURE S2

**Fig. S3 Plant growth measurements. The growth of Brachypodium mutant plants grown under greenhouse conditions was measured every week and was compared to Wild-type.** (a) Brachypodium *xax1a* mutant plants appear dwarfed compared to Wild-type. 17 individual plants were measured for Wild-type and 20 individual plants were measured for *bdxax1a* mutant. (b) Dwarfed phenotype of *bdxax1a* plants at 7 weeks (c) Brachypodium *bdgux2* mutant plants grow similar to Wild-type. (d) *bdgux2* plants grow similar to Wild-type (7 weeks).

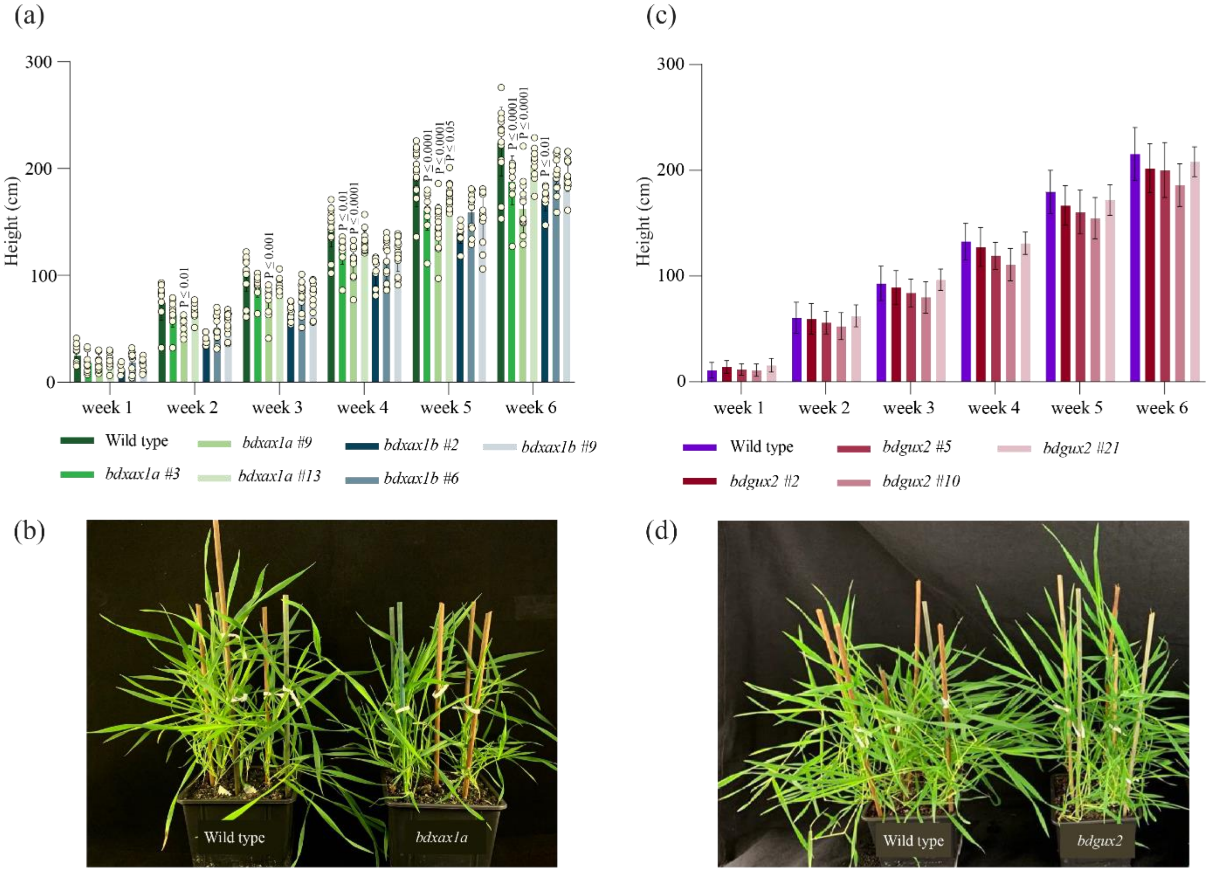


FIGURE S3

**Fig. S4 Gene expression profiles at different Brachypodium developmental stages.** (a) Expression of Bradi1g06560 (*Bd*XAX1A), we chose last internode (young) leaf tissue to perform all biochemical characterisations; (b) Expression of Bradi1g72350 (*Bd*GUX2), we chose last internode tissue to perform all biochemical characterisations.. Expression data were obtained from PlaNet database (http://aranet.mpimp-golm.mpg.de/).

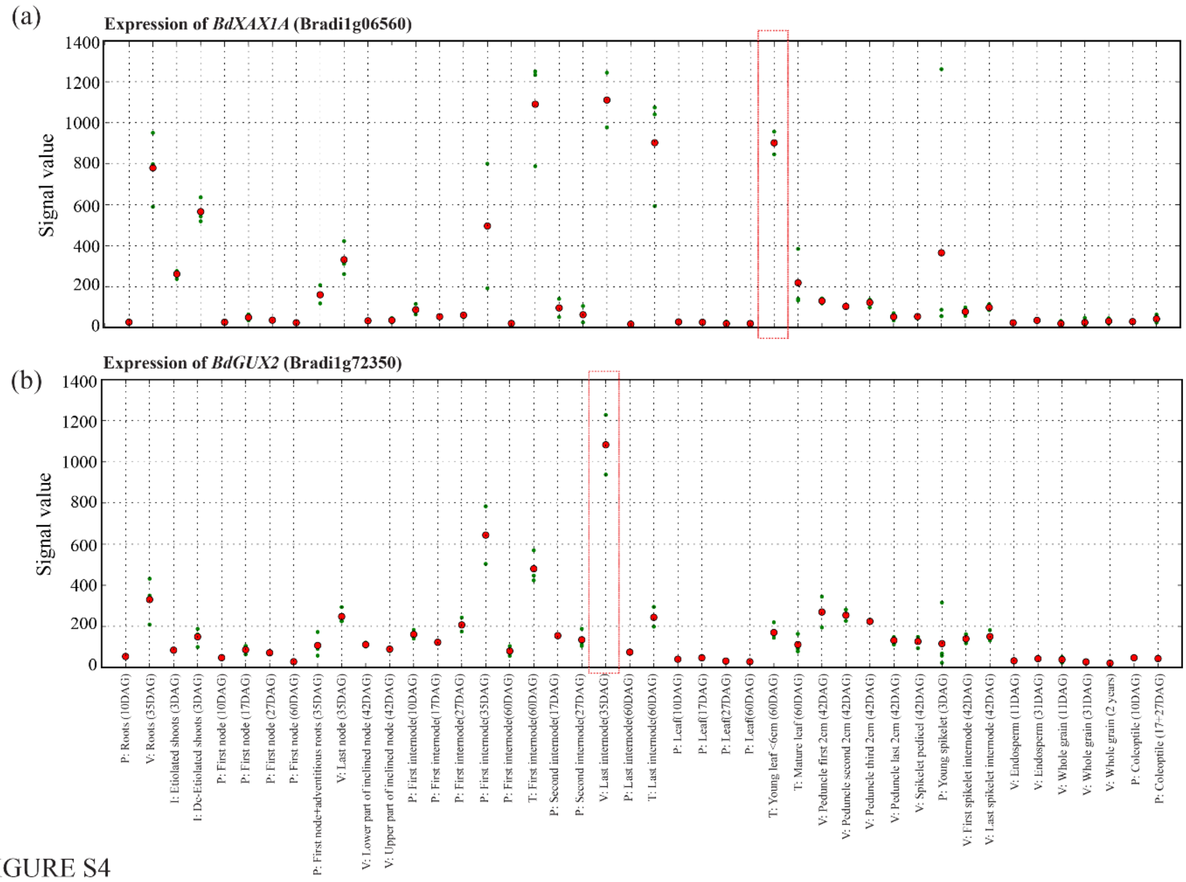
(a)
Expression of BdXAX1A (Bradi1g06560)
Signal value
1400
1200
1000
800
600
400
200
0
(b)
Expression of BdGUX2 (Bradi1g72350)
Signal value
1400
1200
1000
800
600
400
200
0
P: Roots (10DAG)
V: Roots (35DAG)
I: Etiolated shoots (3DAG)
I: De-Etiolated shoots (3DAG)
P: First node (10DAG)
P: First node (17DAG)
P: First node (27DAG)
P: First node (60DAG)
P: First node+adventitious roots (35DAG)
V: Last node (35DAG)
V: Lower part of inclined node (42DAG)
V: Upper part of inclined node (42DAG)
P: First internode(10DAG)
P: First internode(17DAG)
P: First internode(27DAG)
P: First internode(35DAG)
P: First internode(60DAG)
T: First internode(60DAG)
P: Second internode(17DAG)
P: Second internode(27DAG)
V: Last internode(35DAG)
P: Last internode(60DAG)
T: Last internode(60DAG)
P: Leaf(10DAG)
P: Leaf(17DAG)
P: Leaf(27DAG)
P: Leaf(60DAG)
T: Young leaf <6cm (60DAG)
T: Mature leaf (60DAG)
V: Peduncle first 2cm (42DAG)
V: Peduncle second 2cm (42DAG)
V: Peduncle third 2cm (42DAG)
V: Peduncle last 2cm (42DAG)
V: Spikelet pedicel (42DAG)
P: Young spikelet (3DAG)
V: First spikelet internode (42DAG)
V: Last spikelet internode (42DAG)
V: Endosperm (11DAG)
V: Endosperm (31DAG)
V: Whole grain (11DAG)
V: Whole grain (31DAG)
V: Whole grain (2 years)
P: Coleoptile (10DAG)
P: Coleoptile (17+27DAG)

FIGURE S4

**Fig. S5 MALDI-MS analysis of NaOH-sensitive xylan hydrolysis products of 20% ethanol fraction of the SPE $C_{18}$ eluate for Wild-type and *bdxax1a* mutant lines**. Samples were derivatised with procainamide and analysed at reflector positive mode. X, Xyl; A, Ara*f*; FA, ferulic acid; *p*CA, *para*-coumaric acid.

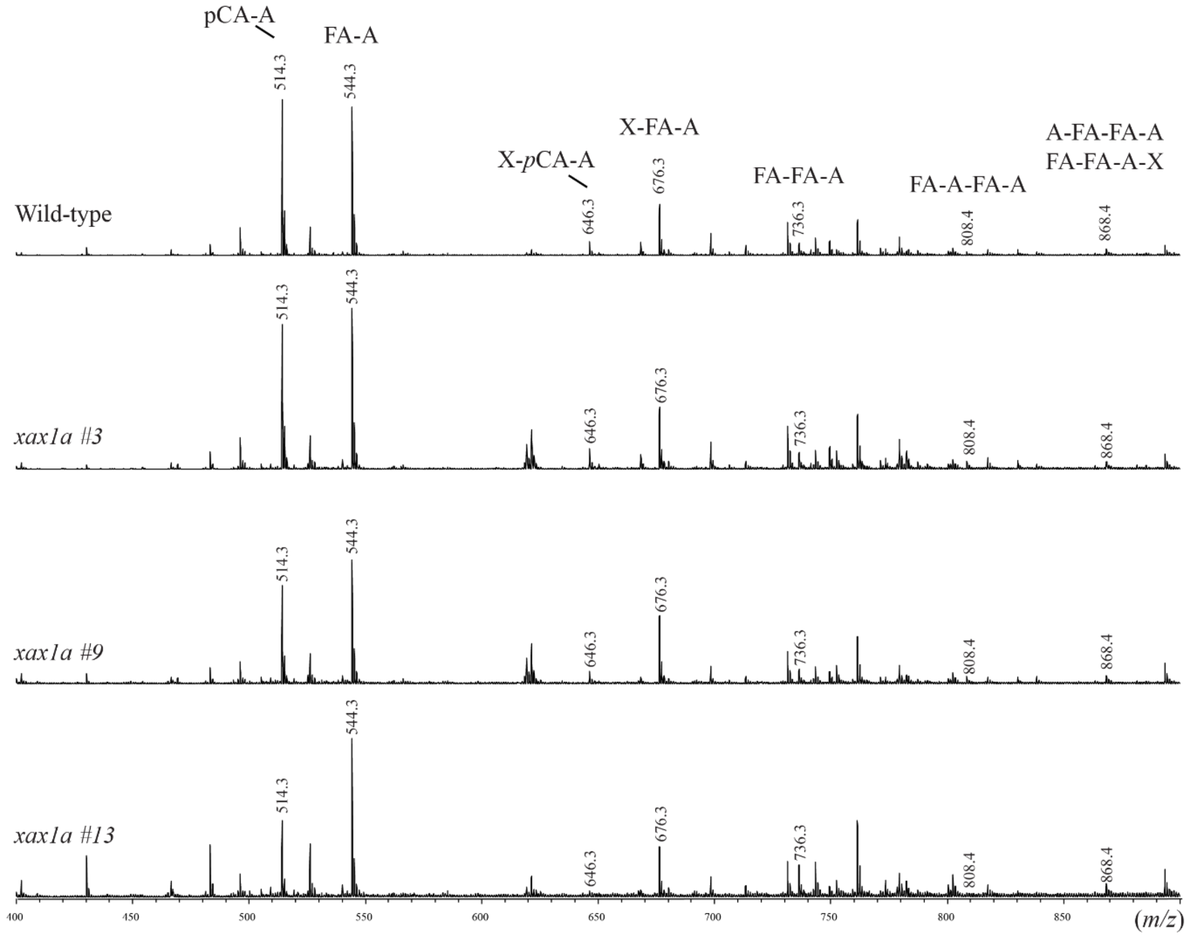


FIGURE S5

**Fig. S6 Sequential cell wall extraction from Brachypodium Wild-type and *bdxax1a* young leaves.** Cell walls were sequentially extracted using CDTA, $Na_2CO_3$, 1M KOH, 4M KOH and 4M KOH post sodium chlorite treatment (4M KOH). Resulting fractions were analysed by PACE using different xylanases. Only, trace amounts of xylan were released by 1,2-cyclohexanediamine tetra acetic acid (CDTA) and $Na_2CO_3$ fractions, which solubilise mainly pectic polysaccharides, from Wild-type walls. (a) A significant amount of GH10 digestible xylan was extracted with $Na_2CO_3$ from *bdxax1a* mutant walls, consistent with Chiniquy *et. al* findings. The majority of (b) AXe (GH5 accessible) and (c) GAXc (digested by the GH30 glucuronoxylanase) were solubilised by 1M KOH, both for Wild-type and *bdxax1a* mutant. A smaller but substantial amount of AXe and trace amounts of GAXc was found in 4M KOH fraction, from both genotypes. Xylan oligosaccharides: X-$X_6$.

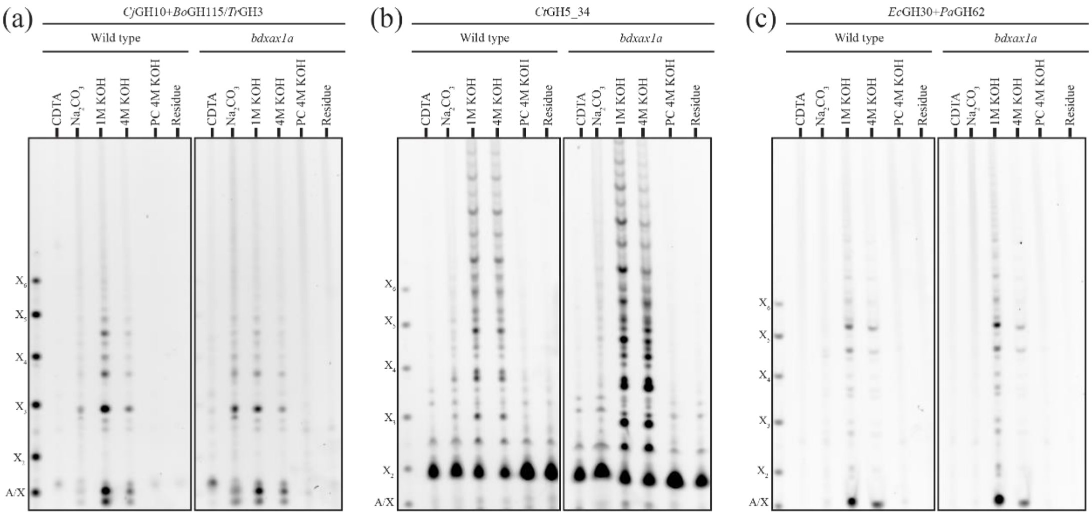


FIGURE S6

**Fig. S7 Characterisation of subcritical water extraction fractions (SWE) from Brachypodium Wild-type and *bdxax1a* young leaves at different timepoints.** To avoid degradation of xylan induced by SWE at high temperatures and acidic pH conditions, a buffered aqueous solution (pH 7.0) was used to minimize degradation. Subcritical water extractions were carried out at 160 °C for 15, 30, 60 and 120 minutes. Monosaccharide and hydroxycinnamate content were monitored across all fractions. (a) Total solids yield (% DW) from Wild-type and *bdxax1a* young leaf samples. Bars represent median and error bars show 95% confidence interval of media for the data; (b) Arabinose to Xylose (Ara:Xyl) ratio of SWE fractions for Wild-type and *bdxax1a* mutant. Error bars represent standard error of three Wild-type and mutant technical replicates. (c) Quantification of SWE fraction total hydroxycinnamates (mg $ml^{-1}$) for Wild-type and *bdxax1a* mutant by HPLC. Error bars represent standard error of three Wild-type and mutant technical replicates. (d) Size-exclusion chromatography (SEC) analysis (d) alkali (4M NaOH) extracted xylan from young leaves of Brachypodium Wild-type and *bdxax1a* mutant; (e) SWE 30 min fraction; (f) SWE 60 min fraction and (g) SWE 120 min fraction. The elugrams represent the signals from the UV detector (in blue) and the refractive index (RI) detector (in orange). Calibration molecular weight standards (pullulan) at corresponding elution times are also shown.

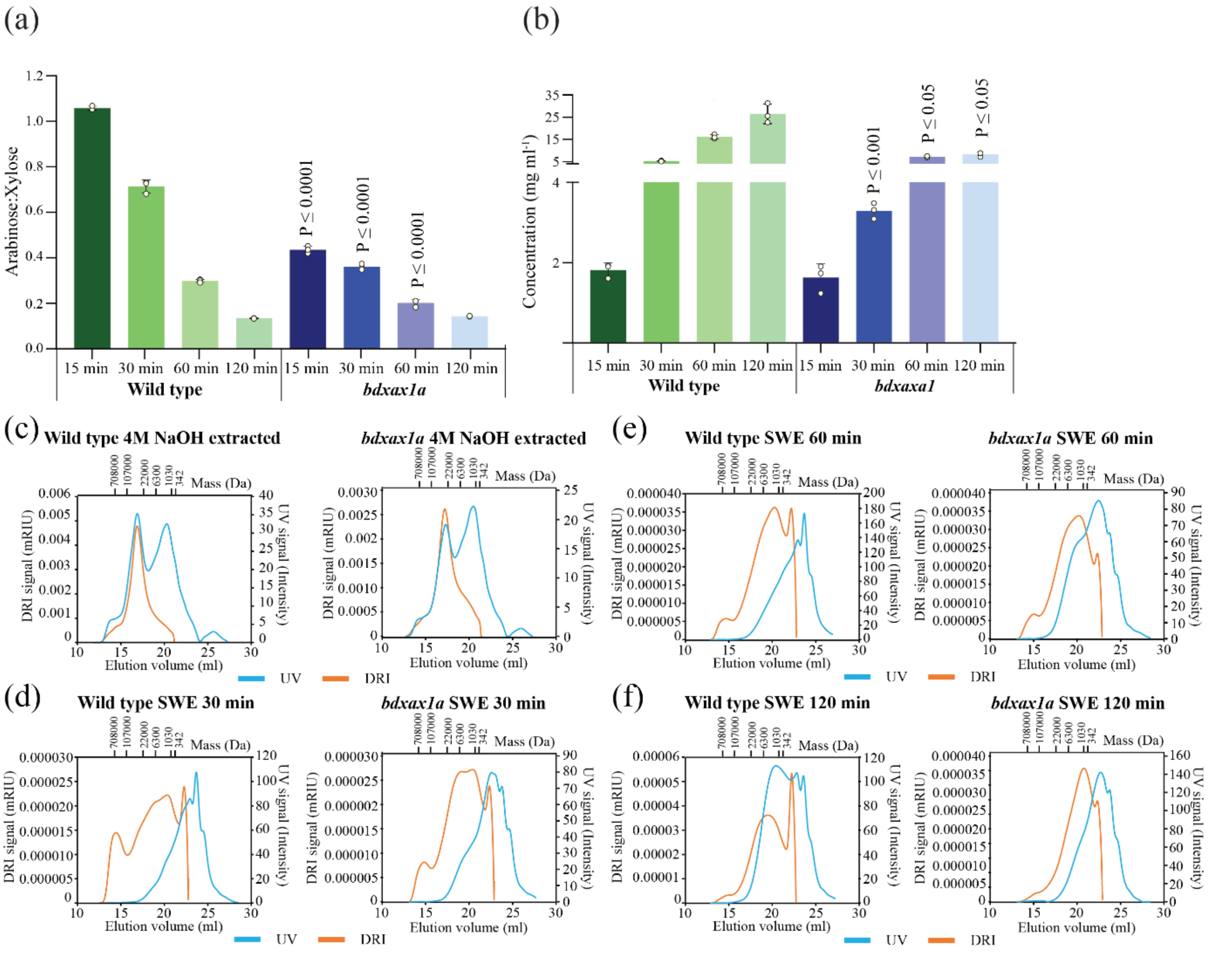

(a)
Arabinose:Xylose
15 min 30 min 60 min 120 min 15 min 30 min 60 min 120 min
Wild type
bdxax1a
P ≤ 0.0001
(b)
Concentration (mg ml-1)
P ≤ 0.001
P ≤ 0.05
Wild type
bdxaxa1
(c) Wild type 4M NaOH extracted
bdxax1a 4M NaOH extracted
(d) Wild type SWE 30 min
bdxax1a SWE 30 min
(e) Wild type SWE 60 min
bdxax1a SWE 60 min
(f) Wild type SWE 120 min
bdxax1a SWE 120 min
Mass (Da)
708000 107000 22000 6300 1030 342
DRI signal (mRIU)
UV signal (Intensity)
Elution volume (ml)
UV
DRI


FIGURE S7

**Fig. S8 Sequential cell wall extraction from Brachypodium Wild-type and *bdgux2* young leaves.** Cell walls were sequentially extracted using CDTA, $Na_2CO_3$, 1M KOH, 4M KOH and 4M KOH post sodium chlorite treatment (4M KOH). Resulting fractions were analysed by PACE using different xylanases. (a) No detectable amounts of xylan were released by CDTA and only trace amounts of xylan were released by $Na_2CO_3$ fractions. The majority of (b) AXe (GH5 accessible) and (c) GAXc (digested by the GH30 glucuronoxylanase) were solubilised by 1M KOH, both for Wild-type and *gux2* mutant, with a small amount of AXe found in 4M KOH fraction. Xylan oligosaccharides: X-$X_6$.

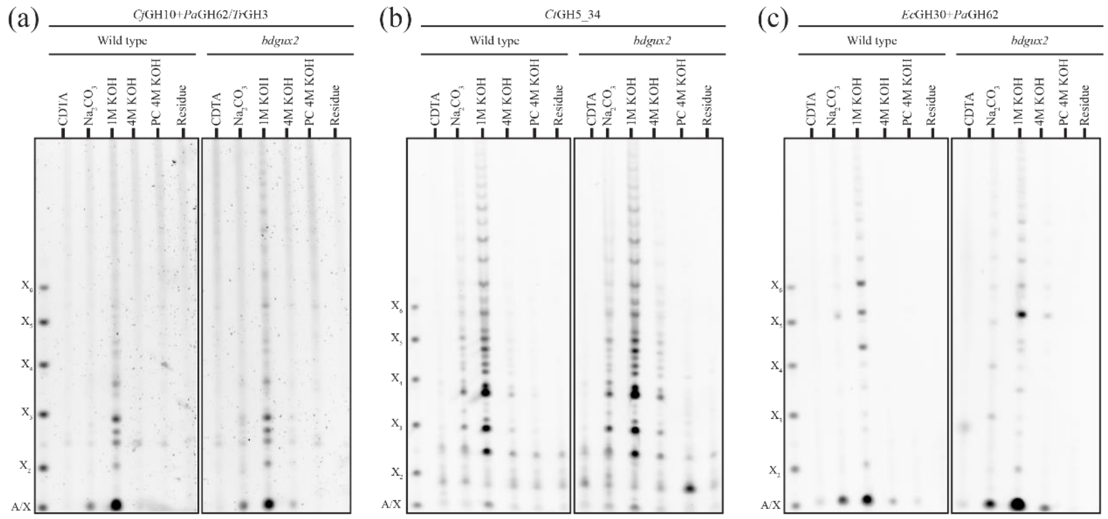


FIGURE S8

**Fig. S9 Characterisation of subcritical water extraction fractions (SWE) from Brachypodium Wild-type and *gux2* last internodes at different timepoints.** To avoid degradation of xylan induced by SWE at high temperatures and acidic pH conditions, a buffered aqueous solution (pH 7.0) was used to minimize degradation. Subcritical water extractions were carried out at 160 °C for 15, 30, 60 and 120 minutes. Monosaccharide and hydroxycinnamate content were monitored across all fractions. (a) Total solids yield (% DW) from wild-type and *gux2* last internode samples. Bars represent median and error bars show 95% confidence interval of media for the data. Size-exclusion chromatography (SEC) analysis (b) alkali (4M NaOH) extracted xylan from last internodes of Brachypodium wild type and *gux2* mutant; (c) SWE 30 min fraction; (d) SWE 60 min fraction and (e) SWE 120 min fraction. The elugrams represent the signals from the UV detector (in blue) and the refractive index (RI) detector (in orange). Calibration molecular weight standards (pullulan) at corresponding elution times are also shown.

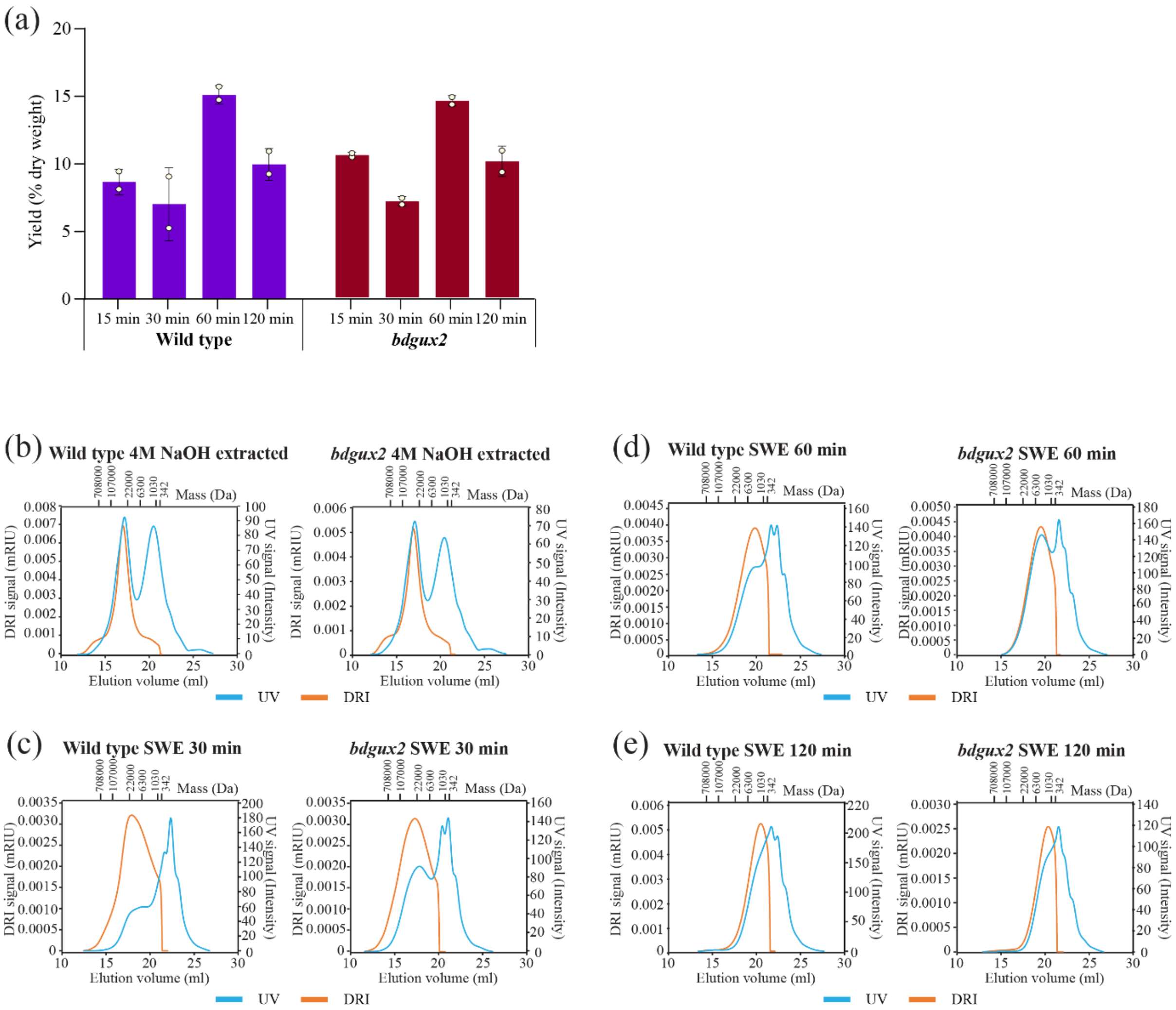
(a)
Yield (% dry weight)
15 min 30 min 60 min 120 min
Wild type
15 min 30 min 60 min 120 min
bdgux2
(b) Wild type 4M NaOH extracted
bdgux2 4M NaOH extracted
(c) Wild type SWE 30 min
bdgux2 SWE 30 min
(d) Wild type SWE 60 min
bdgux2 SWE 60 min
(e) Wild type SWE 120 min
bdgux2 SWE 120 min
708000 107000 22000 6300 1030 342 Mass (Da)
DRI signal (mRIU)
UV signal (Intensity)
Elution volume (ml)
UV DRI

FIGURE S9

**Fig. S10. Comparison of carbohydrate regions of 1D $^{13}C$ NMR spectra among *Brachypodium* samples.** (a) Overlay of young leaf spectra: Wild-type (black) and *bdxax1a* mutant (red). The spectra include (from top to bottom): quantitative DP (q-DP) spectra with 35 s recycle delay, CP spectra, DP spectra with 2 s recycle delay, and INEPT spectra. Assignments of the major cell wall components are indicated for arabinose (A) carbon 1 (C1) and carbon 2 (C2); three-fold xylan ($Xn^{3f}$) C1, C2 and C3; interior cellulose (iC) C4; surface cellulose (sC) C4; two-fold xylan ($Xn^{2f}$) C4; methyl groups (MeO) of ferulate (FA), syringyl (S) and guaiacyl (G) species of lignin. Mutant spectra were simultaneously scaled to match the signal of interior cellulose iC C4 signal at 89 ppm (marked with an asterisk (*)) in Wild-type CP spectra. All spectra are normalised to number of scans, with an additional scaling applied for convenience (the scaling factor is indicated above each spectrum). Insets highlight differences in arabinose A C1 and/or three-fold xylan C1 signal intensities. (b) Overlay of last internode spectra: Brachypodium Wild-type (black) and *bdgux2* mutant (red). The spectra follow same order as in (a), with identical normalization procedure employed.

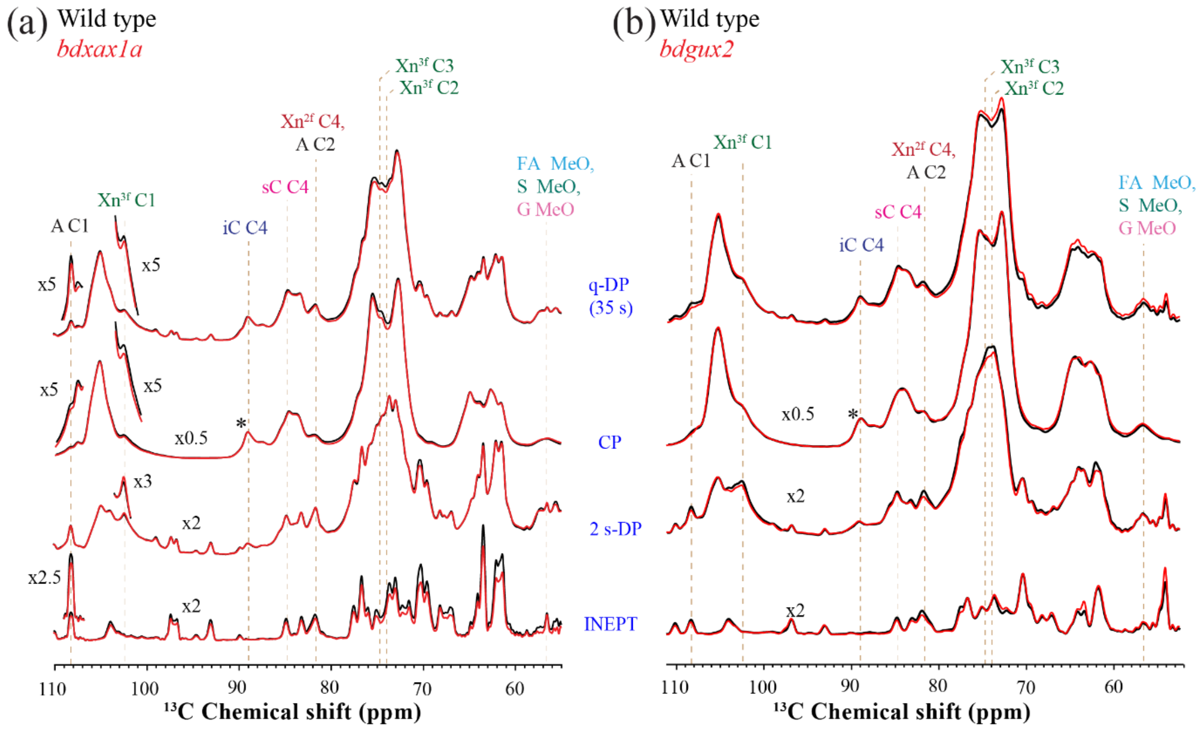
(a) Wild type
bdxax1a
Xn3f C3
Xn3f C2
Xn2f C4,
A C2
sC C4
iC C4
Xn3f C1
A C1
FA MeO,
S MeO,
G MeO
x5
x5
x5
x5
x0.5
*
x3
x2
x2.5
x2
q-DP
(35 s)
CP
2 s-DP
INEPT
110
100
90
80
70
60
13C Chemical shift (ppm)
(b) Wild type
bdgux2
Xn3f C3
Xn3f C2
Xn3f C1
A C1
Xn2f C4,
A C2
sC C4
iC C4
FA MeO,
S MeO,
G MeO
x0.5
*
x2
x2
110
100
90
80
70
60
13C Chemical shift (ppm)

FIGURE S10

**Fig. S11 Full 1D $^{13}C$ NMR spectra of *Brachypodium* young leaf samples.** The spectra of Wild-type (black) and *xax1a* (red) cell walls are overlaid. Spectra from top to bottom are: Q-DP spectra measured with a 35 s recycle delay, 0.5 ms CP spectra, DP spectra measured with a 2 s recycle delay, and INEPT spectra. Mutant spectra were simultaneously scaled to match the signal of the interior cellulose iC C4 at 89 ppm (indicated with an asterisk) in the Wild-type 0.5 ms CP spectra. An additional scaling factor of x 0.5 is applied to the CP spectra for clarity. Insets with 5x intensity magnification highlight the signals of lignin species. *s/b*: spinning sidebands.

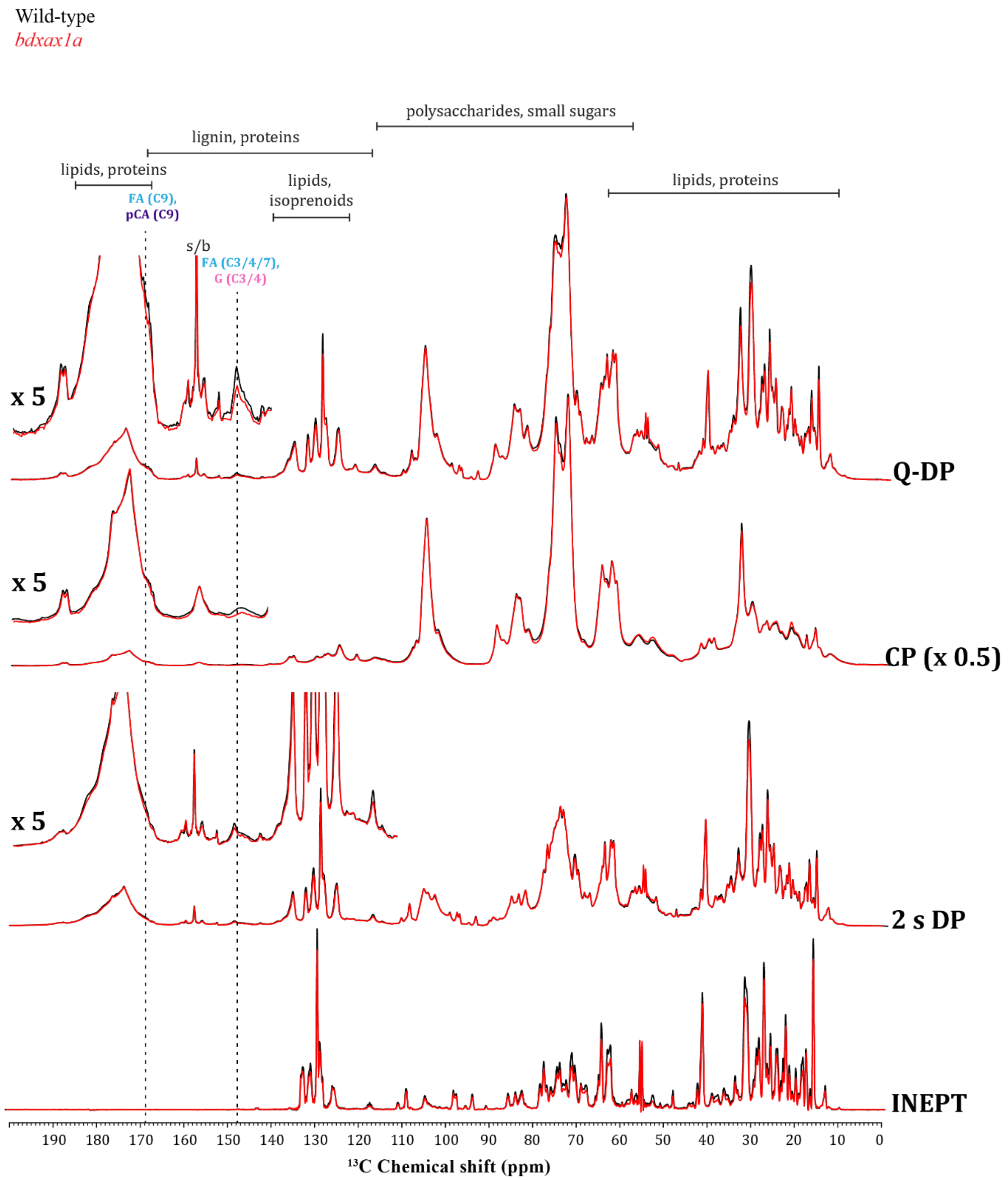


FIGURE S11

**Fig. S12 2D $^{13}$C–$^{13}$C CP-INADEQUATE spectra of *Brachypodium* young leaf cell walls.** The Wild-type (black) and *bdxax1a* (red) young leaf spectra are shown on the left, with integration areas highlighted in yellow. The key cross-sections extracted at the indicated $\omega_1$

double quantum chemical shifts of the Wild-type (black) and *bdxax1a* mutant (red) young leaf cell walls are shown on the right. The computed ratio between the integrated areas of $Xn^{3f}$ C4 cross-peak at (140.5, 77.3) ppm in xax1a and Wild-type spectra as well as the ratio between the integrated areas of $Xn^{2f}$ C4 cross-peak at (146.4, 82.2) ppm in these spectra are shown in the corresponding cross-sections. These integrated areas were normalised by the integrated area of the iC C4 peaks.

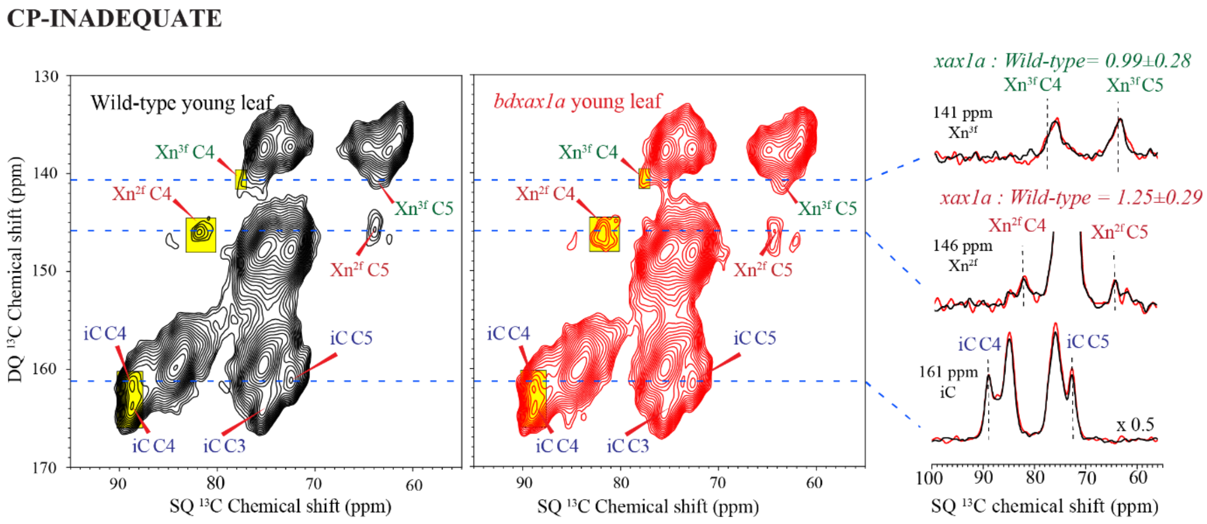


FIGURE S12

**Figure S13. 2D $^{13}$C–$^{1}$H dipolar-doubled CP-DIPSHIFT spectra of *Brachypodium* young leaf and last internode cell walls.** (a) CP-DIPSHIFT dipolar dephasing curves of representative sites of Wild-type (black) and *xax1a* (red) mutant young leaf. The best-fit dipolar order parameters $S_{sample}$ are given in each panel. Intensity uncertainties asre propagated from spectral noise and are < 5% for all sites, which are too small to display. (b) CP-DIPSHIFT dipolar dephasing curves of representative sites of Wild-type (black) and *gux2* (red) mutant last internode cell walls. Error bars represent the uncertainty propagated from spectral noise. Intensity uncertainties are propagated from spectral noise and are < 7% for the $Xn^{3f}$ and $Xn^{2f}$ sites, which are too small to display. (c) Cross sections of the 2D $^{13}$C–$^{1}$H CP-DIPSHIFT spectra without dipolar evolution ($t_{dip}$ = 0 μs) and with maximum dipolar evolution ($t_{dip}$ = 47.62 μs) for *bdxax1a* mutant (red) and Wild-type (black) young leaf samples. (d) Cross sections of the 2D $^{13}$C–$^{1}$H quantitative dipolar-doubled DIPSHIFT spectra without dipolar evolution ($t_{dip}$ = 0 μs) and with maximum evolution ($t_{dip}$ = 57.35 μs) for *gux2* mutant (red) and Wild-type (black) last internode samples.

**0.5 ms CP $^{13}C$-$^{1}H$ DIPSHIFT**

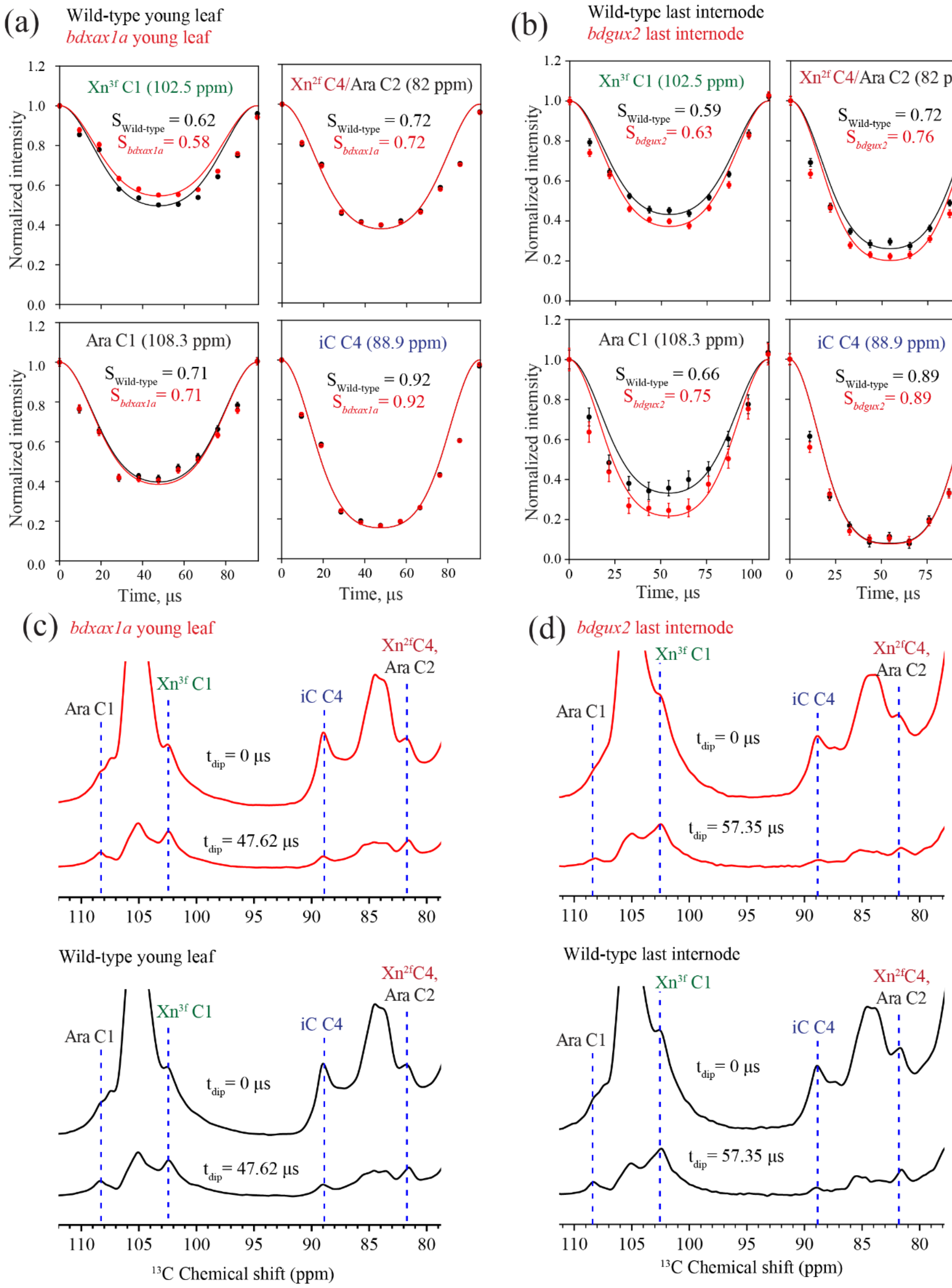


FIGURE S13

**Figure S14. Representative cross sections of the 2D $^{13}C-^{1}H$ dipolar-doubled Q-DP DIPSHIFT spectra of *Brachypodium* young leaf and last internode cell walls.** (a) Cross sections of the 2D q-DP DIPSHIFT spectra without dipolar evolution ($t_{dip}$ = 0 μs) and with maximum evolution ($t_{dip}$ = 47.62 μs) for *xax1a* mutant (red) and Wild-type (black) young leaf samples. (b) Cross sections of the 2D DIPSHIFT spectra without dipolar evolution ($t_{dip}$ = 0 μs) and with maximum evolution ($t_{dip}$ = 57.35 μs) for *gux2* mutant (red) and Wild-type (black) last internode samples.

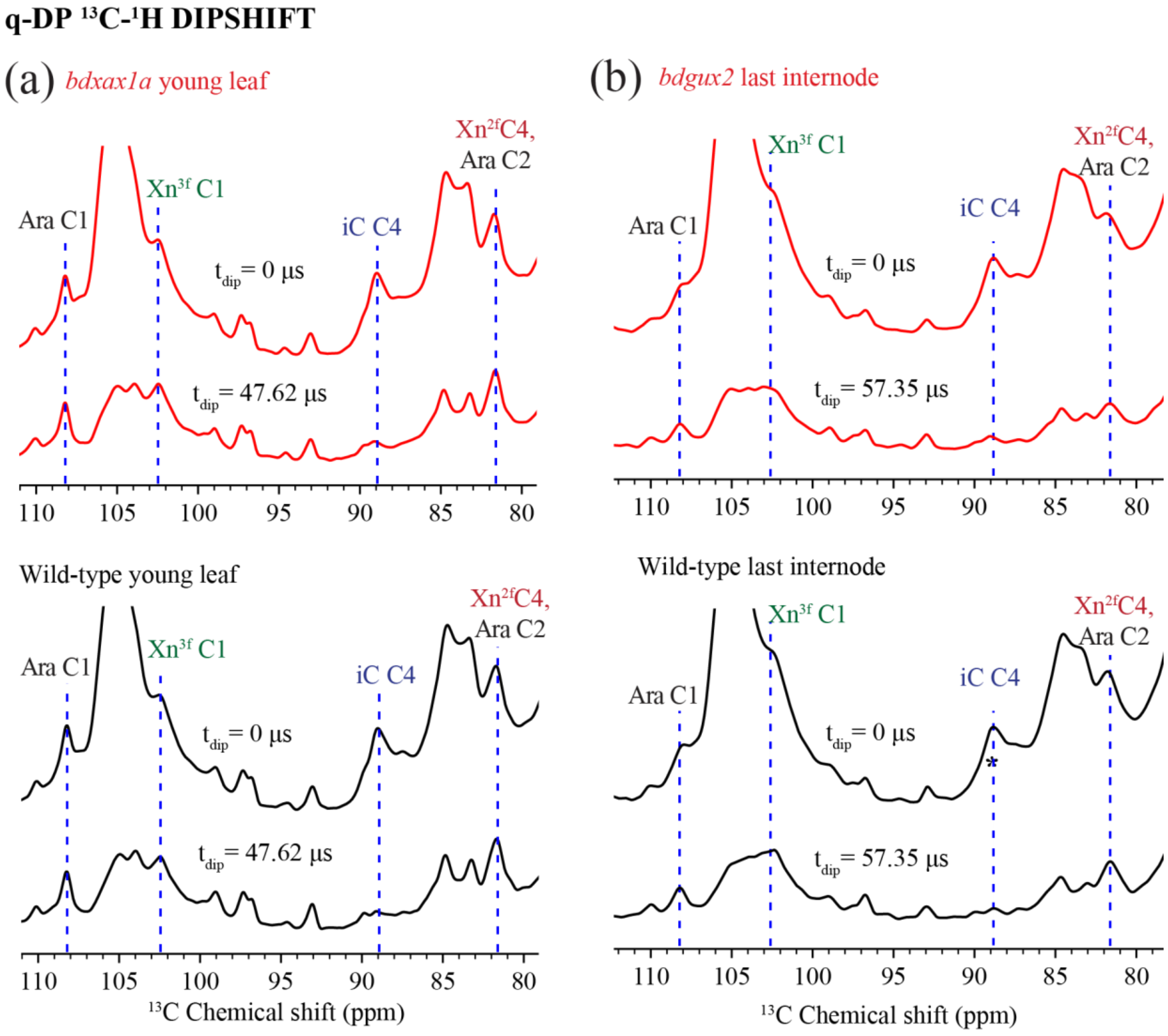


FIGURE S14

**Supplementary Tables**

**Table S1.** Summary of genotypes of CRISPR Cas9 lines. Homozygous or bi-allelic (two different mutations occurred on each allele of the targeted gene) transgenic lines were regenerated, and 3-4 transgenic lines were biochemically analyzed as later described. Transgenic lines were genotyped by PCR amplifying amplicons spanning the DSB site, using the marked primers flanking the DSB site. The PCR products were then purified and sequenced to determine the INDELS which occurred at the DSB sites in comparison to also sequenced Wild-type control PCR amplicons amplified using the same primer pairs per gene. The GUX2 CRISPR transgenic lines 2, 10, and 21 were determined to be bi-allelic for 1 nucleotide insertions on each allele versus 2 nucleotides inserted on one allele, and the other allele being of Wild-type sequence, based on the fact that all heterozygous lines which were also genotyped but not biochemically analyzed were of normal height, rather than stunted. This indicated that there was not a Wild-type allele remaining in the stunted GUX2 lines 2, 10, and 21 to confer normal height, and rather that the 2 nucleotides inserted must be on opposite alleles.

| Construct | Line # | Genotype | Mutation(s) |
|---|---|---|---|
| BdXAX1a | 3 | Homozygous | Deletion of 1 (T) on both alleles |
| BdXAX1a | 9 | Homozygous | Deletion of 1 (T) on both alleles |
| BdXAX1a | 13 | Homozygous | Deletion of 17 nucleotides in combination with insertion of 4 nucleotides |
| BdXAX1b | 2 | Homozygous | Insertion of 1 (A) on both alleles |
| BdXAX1b | 6 | Homozygous | Deletion of 1 (G) on both alleles |
| BdXAX1b | 9 | Homozygous | Insertion of 1 (A) on both alleles |
| BdGUX2 | 2 | Bi-allelic | Insertion of 1 (A) on one allele; Insertion of 1 (T) on other allele |
| BdGUX2 | 5 | Homozygous | Insertion of 1 (T) on both alleles |
| BdGUX2 | 10 | Bi-allelic | Insertion of 1 (A) on one allele; Insertion of 1 (T) on other allele |
| BdGUX2 | 21 | Homozygous | Insertion of 1 (T) on both alleles |

**Table S2.** List of Primers used in this study. Sequences are shown from 5’ to 3’.

| Primer Name | Primer Description | Sequence (5’ to 3’) |
|---|---|---|
| pCTA2390 | eGFP exog CRISPR for Bd F1 no PAM | ggcaTGAGCAAGGGCGAGGAGCTG |
| pCTA2391 | eGFP exog CRISPR for Bd R1 no PAM | aaacCAGCTCCTCGCCCTTGCTCA |
| pCTA2826 | Bradi1g06560 XAX1a gRNA1-F | ggcaGGAGCCCGAGATGAATTCCA |
| pCTA2827 | Bradi1g06560 XAX1a gRNA1-R | aaacTGGAATTCATCTCGGGCTCC |
| pCTA2828 | Bradi3g11337 XAX1b gRNA1-F | ggcaGCGAGAGGAGGCTCCCGCCG |
| pCTA2829 | Bradi3g11337 XAX1b gRNA1-R | aaacCGGCGGGAGCCTCCTCTCGC |
| pCTA2572 | BdGUX2 gRNA1-F - targeting Bradi1g72350 | taggtctcaTCATCAAGTCTCgttttagagctagaa |
| pCTA2573 | BdGUX2 gRNA1-R - targeting Bradi1g72350 | cgggtctcaATGAGCGACTCAtgcaccagccggg |
| pCTA2944 | BdXAX1a genotyping F3 (Bradi1g06560) | GGTTGCTTCGAGCAGATAAA |
| pCTA2946 | BdXAX1a genotyping R3 (Bradi1g06560) | GTTGAGTACGCACCGAATAC |
| pCTA2947 | BdXAX1b genotyping F3 (Bradi3g11337) | CTGCTCATTTCAGTAGCCAATTAC |
| pCTA2948 | BdXAX1b genotyping R3 (Bradi3g11337) | GCTTCTTGGGCTCGATCTT |
| pCTA2940 | BdGUX2 F genotyping primer (targeting Bradi1g72350) | GAGACCGTGAGACACAACAA |
| pCTA2941 | BdGUX2 R genotyping primer (targeting Bradi1g72350) | GCACCTTACCTTCCGTGAG |

**Table S3. Experimental conditions for the $^{13}$C MAS NMR.**

The choice of spectrometer $^{1}$H frequency was dictated only by the availability of the spectrometer at the moment of data collection. All experiments were done at sample temperature 278 K on an 800 MHz (18.8 T) Bruker Avance II or a 600 MHz (14.1 T) Bruker Avance II spectrometer, using HCN probe in double resonance mode. Typical radiofrequency (rf) field strengths were 56.8 kHz for $^{13}$C and 71.4 kHz for $^{1}$H. Proton decoupling was conducted using the two-pulse phase-modulation (TPPM) (Bennett *et al.*, 1995) scheme with an rf field strength of 71.4 kHz.

| Sample | Experiment | $^{1}$H Larmor frequency, MAS rate | Mixing time (if used), CP contact time (if used), recycle delays (D1), Number of scans (NS), Digitization |
|---|---|---|---|
| Young leaves, *bdxax1a* and Wild-type | 1D $^{13}$C quantitative DP | 800 MHz, 10.5 kHz | D1 35 s, NS 1024 |
| | 1D $^{13}$C CP | | 500 µs $^{1}$H-$^{13}$C CP, D1 1.7 s, NS 2048 |
| | 1D $^{13}$C DP with short recycle delay | | D1 2 s, NS 2048 |
| | 1D $^{13}$C INEPT | | D1 1.7 s, NS 2048 |
| | 2D $^{13}$C–$^{1}$H quantitative DP dipolar-doubled DIPSHIFT | | D1 25 s, NS 80, TD2 2560, TD1 11, FSLG 83 kHz |
| | 2D $^{13}$C–$^{1}$H CP dipolar-doubled DIPSHIFT | | 500 µs $^{1}$H-$^{13}$C CP contact time, D1 1.7 s, NS 320, TD2 2560, TD1 11; FSLG 83 kHz |
| | 2D $^{13}$C-$^{13}$C double-quantum and single-quantum CP-INADEQUATE | 600 MHz, 9.8 kHz | 500 µs $^{1}$H-$^{13}$C CP; SPC-5 recoupling 49 kHz ($5w_r$) ; NS 240, TD2 1600, TD1 292, IN1 34 µs |
| | 1D water-edited $^{13}$C CP | 600 MHz, 11.4 kHz | $^{1}$H spin diffusion times (ms): 1; 2; 4; 9; 16; 25; 36; 49; 100. Gaussian $^{1}$H 180˚ pulse: 1.4 ms; 500 µs $^{1}$H-$^{13}$C CP; NS 5120 |
| Last internode, *bdgux2* and Wild-type | 1D $^{13}$C quantitative DP | 600 MHz, 11.4 kHz | D1 35 s, NS 512 |
| | 1D $^{13}$C CP | | 500 µs $^{1}$H-$^{13}$C CP contact time, D1 2 s, NS 1024 |
| | 1D $^{13}$C DP with short recycle delay | | D1 2 s, NS 2048 |
| | 1D $^{13}$C INEPT | | D1 2 s, NS 2048 |
| | 2D $^{13}$C–$^{1}$H quantitative DP dipolar-doubled DIPSHIFT | 600 MHz, 9.2 kHz | D1 25 s, NS 192, TD2 2048, TD1 11, FSLG 62.5 kHz |
| | 2D $^{13}$C–$^{1}$H CP dipolar-doubled DIPSHIFT | | 500 µs $^{1}$H-$^{13}$C CP contact time, D1 1.7 s, NS 320, TD2 2048, TD1 11; FSLG 62.5 kHz |
| | 1D water-edited $^{13}$C CP | 600 MHz, 11.4 kHz | $^{1}$H spin diffusion times (ms): 2; 4; 9; 16; 25; 49; 100; 225. Gaussian $^{1}$H 180˚ pulse duration 1.4 ms; NS 2048 |

**Table S4. $^{13}$C Chemical shifts of major polysaccharide and lignin species.**

The tentative chemical shifts assignment is based on literature chemical shifts (Simmons *et al.*, 2016; Kang *et al.*, 2019).

| Species | $^{13}$C Chemical shift, ppm |
|---|---|
| $Xn^{3f}$: C1; C2; C3 | 102.4; 73.7; 74.5 |
| $Xn^{2f}$: C4 | 81.6 |
| Ara: C1; C2 | 108.3; 81.6 |
| iC: C4 | 88.9 |
| sC: C4 | 83.7 |
| FA: C3, C4, C9, OMe | 148; 148; 169; 56.7 |
| *p*CA: C9 | 169 |
| G: C3, C4, OMe | 148; 148; 56.7 |
| S: OMe | 56.7 |

**Supplementary Methods**

**Methods S1.** Sold-state NMR experiments.

Approximately 40 mg of each never-dried [$^{13}$C]-labeled *Brachypodium bdxax1a* young leaf, Wild-type young leaf, *bdgux2* last internode, and Wild-type last internode tissues were separately packed into Bruker thin-wall 3.2-mm MAS rotors. To prevent sample degradation, approximately 6 µl of 20 mM $NaN_3$ solution was added to each packed sample, and sample temperatures were maintained at ~278 K during NMR measurements. Packed samples were stored at -20 ºC between measurements. Sample temperature $T_{sample}$ (measured in Kelvin) during NMR measurements was determined from $^1$H chemical shift of the water peak $\delta(^1H_2O)$ (measured in ppm, internally calibrated to DSS) according to the equation $T_{sample} = 7.83 - \delta(^1H_2O)/96.9$ (Hartel *et al.*, 1982). $^{13}$C-chemical shifts were externally referenced to the adamantane methylene peak at 38.48 ppm on the tetramethylsilane.

Four types of one-dimensional (1D) $^{13}$C NMR spectra were measured for all the samples: a quantitative direct polarization (DP) experiment that detects all species regardless of mobility; another $^{13}$C DP experiment with a short recycle delay of 2 s that allows to suppress the signals of rigid molecules with slow $T_1$ relaxation while enhancing the signals of dynamic molecules with fast $^{13}$C $T_1$ relaxation; a $^{13}$C refocused insensitive nuclei enhancement by polarisation transfer (INEPT) experiment, which preferentially detect highly dynamic molecules; and a cross-polarization (CP) experiment to selectively detect rigid molecules such as cellulose.

To resolve the signals of immobilised two-fold and three-fold xylan in xax1a and Wild-type young leaf, 2D $^{13}$C- $^{13}$C dipolar-based double-quantum and single-quantum correlation CP-INADEQUATE spectra with the SPC-5 sequence to recouple $^{13}$C-$^{13}$C dipolar coupling under MAS (Hohwy *et al.*, 1999) . This experiment selectively detects rigid polysaccharides.

To investigate polysaccharide dynamics for all the samples we measured dipolar-doubled 2D $^{1-}$H-$^{13}$C dipolar chemical-shift (DIPSHIFT) correlation spectra, using the FSLG sequence for $^{1}$H-$^{1}$H homonuclear decoupling (Munowitz *et al.*, 1981; Hong *et al.*, 1997). Both quantitative direct polarization DP-DIPSHIFT and cross-polarisation CP-DIPSHIFT experiments were conducted. The DIPSHIFT dephasing curves were fitted using a custom-written Python script to obtain the dipolar coupling strengths (available on http://meihonglab.com/hong-lab-software/). The fit values were divided by 22.5 kHz, the rigid-limit one-bond C-H dipolar coupling, and by additional factor of (2 × 0.577) to take into account the dipolar doubling and the scaling factor of FSLG to obtain the order parameter $S_{CH}$.

In 1D water-edited $^{13}$C CP experiments applied to assess water accessibility for all the samples, water $^{1}$H magnetization was selected using selective Hahn echo and transferred to rigid biopolymers through $^{1}$H spin diffusion with variable mixing times (specified in Table S3), followed by $^{1}$H-$^{13}$C CP for $^{13}$C detection (White *et al.*, 2014). To obtain the water-polysaccharide $^{1}$H magnetization transfer buildup curves, we plotted the signal intensity of selected sites as a function of square root of $^{1}$H spin diffusion time. For the resulting curves, we applied a correction for $^{1}$H spin-lattice relaxation ($T_1$), using the equation $I_{corrected}^{i} = I_{experimental}^{i} \cdot \exp(t_{SD}/T_1{}^{i})$, where $I_{experimental}^{i}$ is the intensity of the i$^{th}$ site observed at $^{1}$H spin diffusion mixing time $t_{SD}$, $I_{corrected}^{i}$ is the intensity of the i-th site corrected by $T_1^{i}$ relaxation rate at site i. The values of $T_1^{i}$ were measured via inversion recovery experiments (Carr & Purcell, 1954) (data not shown). To extract the $T_1^{i}$ for each site, we fitted the experimental curve of intensity at chemical shift for site i ($I_{invrec}^{i}$) as a function of $^{1}$H inversion recovery time ($t_{invrec}$) using the equation $I_{invrec}^{i} = 1 - I_0^{i} \cdot \exp(-t_{invrec}/T_1^{i})$ . The parameters $I_0^{i}$ and $T_1^{i}$ for each site were determined through least-squares fitting using custom-written Python script.
